# Epithelial-Mesenchymal Plasticity is regulated by inflammatory signalling networks coupled to cell morphology

**DOI:** 10.1101/689737

**Authors:** Mar Arias Garcia, Zheng Yin, Theodoros I. Roumeliotis, Francesca Butera, Lin Wang, Rebecca Rickman, Jyoti Choudhary, Stephen T.C. Wong, Yinyin Yuan, Chris Bakal

**Affiliations:** Dynamical Cell Systems Team. Chester Beatty Laboratories, The Institute of Cancer Research, 237 Fulham Road, London UK, SW3 6JB; Department of Systems Medicine and Bioengineering, Houston Methodist Cancer Center and Departments of Radiology, Neurosciences, Pathology and Laboratory Medicine, Weill Cornell Medicine, 6670 Bertner Avenue, R6 South, Houston TX, 77030, USA; Functional Proteomics Team. Chester Beatty Laboratories, The Institute of Cancer Research, 237 Fulham Road, London UK, SW3 6JB; Computational Pathology and Integrative Genomics Team. The Institute of Cancer Research 15 Cotswold Road, Sutton, SM2 5NG

## Abstract

Morphology dictates how cells sense physical and soluble cues in their environment; thus contributing to fate decisions. The differentiation of epithelial cells into mesenchymal forms, or epithelial-mesenchymal plasticity (EMP), is essential for metazoan development and homeostasis. Here we show that the decision to engage EMP is coupled to cell morphology by cell-cell adhesions by microtubule and nuclear organization (MTNO). Using an integrative ‘omic approach we identify Junctional Adhesion Molecule 3 (JAM3) as a new tumour suppressor in breast cancer patients. JAM3 depletion in epithelial cells alters MTNO and causes differentiation into mesenchymal forms. Soluble TGFβ also changes MTNO, and synergizes with JAM3 depletion to promote mesenchymal morphogenesis. Through systematic proteomic analysis we show that changes in MTNO lead to the upregulation of an inflammatory signalling network where YAP/TAZ, FOXO, IKK-NFKB, and JNK pathways are active; but where insulin signalling is suppressed. The actions of the MT-motor Kinesin-1 serve to both change MTNO and promote the upregulation of the core EMP network. Critically, the upregulation of the EMP network predicts the mesenchymal state across cancers.

**Graphical Abstract:** 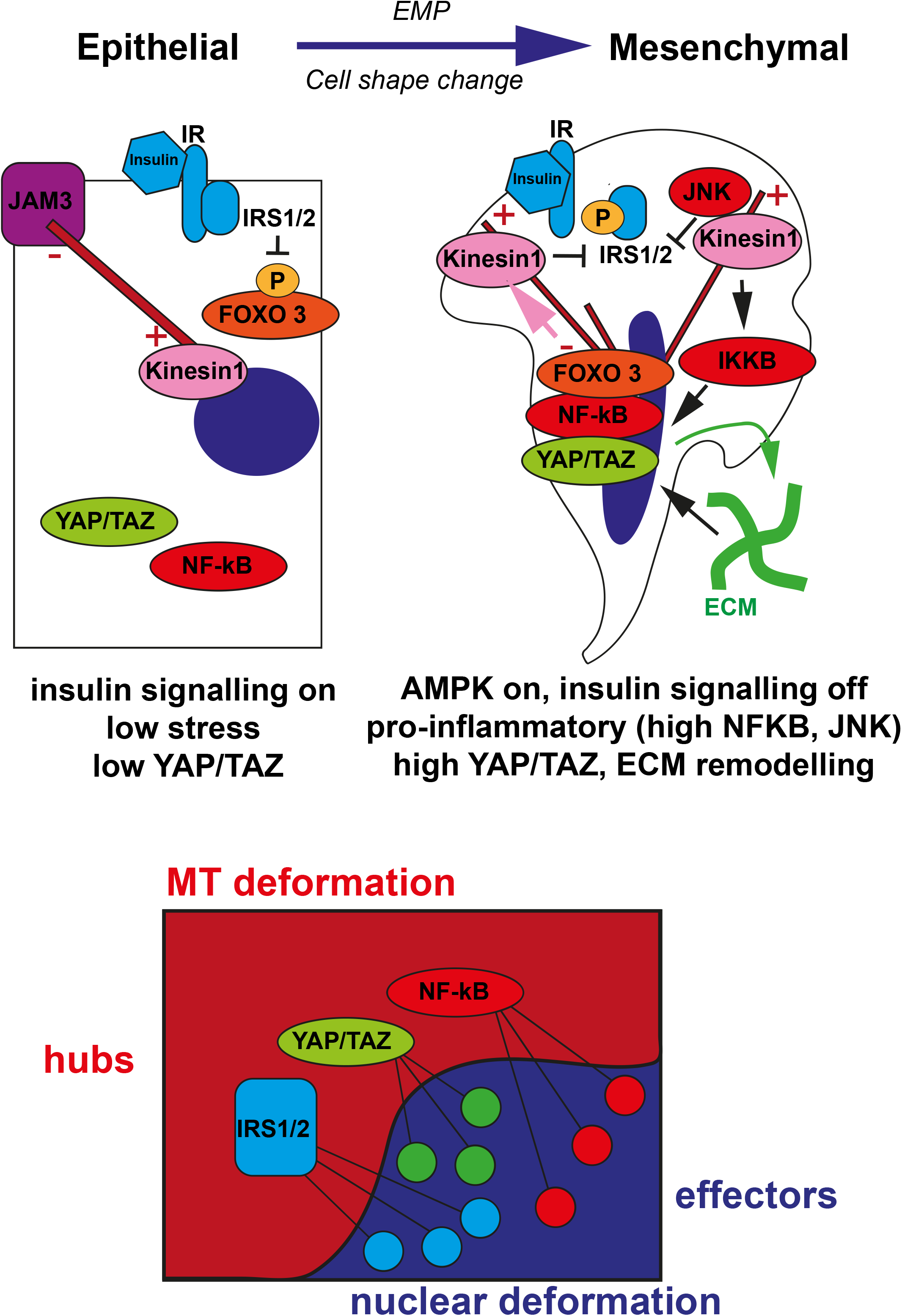

## Introduction

The ability of epithelial cells to differentiate into mesenchymal forms, or epithelial-to-mesenchymal plasticity (EMP) is essential for the development of multiple tissues and homeostatic processes such as wound healing and tissue repair (Nieto et al., 2016; Yang et al., 2021). The difference between epithelial and mesenchymal cells was classically defined by changes in cell shape (Yang et al., 2021). During differentiation from epithelial to mesenchymal forms, cells change from apicobasally polarized epithelial cuboidal cells with cell-cell adhesions such as adherens and tight junctions, to protrusive mesenchymal, fibroblast-type cells with leading edge polarity and the formation of extensive cell-ECM (Extracellular Matrix) adhesions. Dysregulation of EMP has been implicated in cancers, fibrotic diseases, aging related pathologies, and correlates with the severity of infection-driven sequelae such as those following COVID-19 infection (Lambert and Weinberg, 2021; Nieto et al., 2016; Pandolfi et al., 2021).

Transitions from epithelial to mesenchymal forms have been thought to be largely driven by activation of canonical transcription factors such as Slug/SNAI2, Snail, and Twist1 (Batlle et al., 2000; Cano et al., 2000; Leptin, 1991; Nieto et al., 1994; Savagner et al., 1997). But there is a growing list of factors that are implicated in both the engagement and inhibition of EMP. For example, transcriptional activity by NFKB (Sero et al., 2015), YAP/TAZ (Kim et al., 2019; Lehmann et al., 2016; Lei et al., 2008; Seo et al., 2016; Shao et al., 2014; Tang et al., 2016), STAT3 (Saitoh et al., 2016), and FOXO1 (Shin et al., 2019) has been found to promote EMP. While clearly differences in cell shape are important aspect to EMP, the change in signalling state that coincides with EMP also plays a role in development, homeostasis, and disease. For example, mesenchymal states in cancer cells are associated with acquisition of stem cell traits, elevated drug resistance, and increased metastasis (Dongre and Weinberg, 2019; Lambert and Weinberg, 2021). Many of these behaviours are likely driven by the regulatory signalling and transcription factors which themselves drive EMP.

Transitions from epithelial-like to mesenchymal states are triggered by mechanical cues that affect cell shape such as physical disruption of cell-cell adhesion (i.e. wounding), changes in matrix elasticity and ECM geometry (Gomez et al., 2010; Levental et al., 2009; Paszek et al., 2005; Zhan et al., 2008). For example, classic work by Greenburg and Hay demonstrated that suspending single epithelial cells in 3D collagen is sufficient to promote EMP (Greenburg and Hay, 1982). EMT is also be triggered by soluble cues such as TGFβ, FGF, MET, and pro-inflammatory TNFα (Nieto et al., 2016). But how mechanical and soluble cues are coupled to the dynamics of signalling pathways that regulate transcriptional and post-transcriptional events that underpin EMT is very poorly understood. Especially with regards to mechanical cues, it is largely a ‘black box’ as to how changes in adhesion, ECM stiffness, and environment geometry are coupled to the transcriptional events that drive EMP.

We have previously shown that cell and nuclear shape are important determinants of environment affects transcriptional events (Sailem and Bakal, 2017; Sero and Bakal, 2017; Sero et al., 2015). The signalling state can then determine the extent of phenotypic plasticity. For example, in breast cancer cells we showed that epithelial morphologies suppress NFĸB signalling downstream of inflammatory stimuli such as TNFα and prevent the adoption of mesenchymal forms (Sero et al., 2015). The ability of cell and nuclear shape to regulate signalling establishes a means to restrain EMP during inflammation in non-wounded cells (i.e. in cells not at the edge of a wound). However, when stimulated by TNFα the shape of cells in a ‘pre-mesenchymal’ state (i.e. at the edge of wounded monolayers) can act to sustain NFĸB activation and drive EMP through positive feedback (Sero et al., 2015). We have also found that cell and nuclear shapes also control the activity of other transcription factors such as YAP/TAZ (Sero and Bakal, 2017), whose actions can also promote EMP (Kim et al., 2019; Lehmann et al., 2016; Lei et al., 2008; Seo et al., 2016; Shao et al., 2014; Tang et al., 2016). Mechanistically, cell and nuclear morphology can control state in a number of ways. For example, nuclear shape dictates the biophysics of key cellular processes such as transcription, replication, mRNA export, and DNA damage repair (Dahl et al., 2008; Elosegui-Artola et al., 2017; Fedorchak and Lammerding, 2016; Irianto et al., 2016; Irianto et al., 2017; Swift and Discher, 2014).

That nuclear shape in particular as a regulator of cell fate decisions is highlighted by the fact that changes in nuclear morphology and genome organization are directly causal to oncogenesis (Irianto et al., 2017; Pfeifer et al., 2017). Indeed, pathologists have used nuclear shape as a diagnostic for over a century (Chow et al., 2012). Irregular nuclear shapes and sizes are hallmarks of cancer cells, whereas smooth, roundish forms are characteristic of normal cells (Bussolati, 2008). Both the cause and consequences of these nuclear shape deformations in cancer progression are not clear. In some cancer cells, a highly deformable nucleus is necessary to squeeze through small pores (<10 microns) in 3D environments during confined migration (Krause and Wolf, 2015; Pfeifer et al., 2019). Dysregulation in nuclear morphogenesis is also linked to genome instability (Takaki et al., 2017), alterations in the dynamics of DNA damage repair (Irianto et al., 2017; Xia et al., 2018; Xia et al., 2019), and the activation of pro-inflammatory pathways such as canonical and non-canonical NFĸB activation (Bakhoum et al., 2018; Sero et al., 2015) – all of which can contribute to tumorigenesis and metastasis.

Microtubules (MTs) regulate nuclear shape position and structure during flux in environmental and internal conditions (Haase et al., 2016; Wang et al., 2015; Wang et al., 2018). Indeed, MT and nuclear organization (MTNO) are often highly coordinated. Perhaps the best example of which is the separation of chromatin during cell division by the mitotic spindle (Forth and Kapoor, 2017). In epithelial cells, the anchoring of MT minus-ends at cell-cell adhesion complexes results in the stabilization, nucleation, and polymerization of MTs along the apicobasal axis (Akhmanova and Hoogenraad, 2015; Bartolini and Gundersen, 2006; Keating and Borisy, 1999; Meads and Schroer, 1995; Shtutman et al., 2008). Nesprin/SYNE proteins serve as adaptors which link the nuclear membrane to specific cytoskeletal components (Ketema and Sonnenberg, 2011; Rajgor and Shanahan, 2013). The heavy chain of the Kinesin-1 MT-bound motor KIF5B forms a complex with SYNE4 which can determine the positioning of MTs with respect to the nucleus (Roux et al., 2009; Wu et al., 2018). Moreover, MTs, Nesprin-4, and Kinesin-1 are required for the mobility of damaged DSBs (Lottersberger et al., 2015), and the LINC complex required for dynamic movement of chromosomal loci such as telomeres during bouquet formation (Shibuya and Watanabe, 2014).

Because cancer cells have dysregulated nuclear shapes, we reasoned gene expression changes in these cells may point to mechanisms as to how cells either protect or deform nuclei. Thus we first used an integrative omic approach to identify mRNAs whose expression is associated with nuclear shape changes in breast cancer patients. Our analysis revealed that the gene encoding the cell-cell adhesion component JAM3 is a tumour suppressor that maintains nuclear shape *in vivo*. JAM3 is required for maintenance of the epithelial state and its depletion synergizes with canonical EMT-promoter TGFβ (Miettinen et al., 1994) to promote epithelial-mesenchymal plasticity (EMP). Indeed, depletion of JAM3 results in aberrant 3D invasion, defects in organoid morphogenesis and activates YAP/TAZ. EMP following TGFβ stimulation and/or JAM3 depletion can be reversed by: 1) genetic depletion of KIF5B and SYNE-4; 2) by chemical inhibition of Kinesin-1; 3) depletion of TAZ. Thus, we propose Kinesin-1 activity on MTs is essential for EMP.

To understand how Kinesin-1 activity drives shape and state plasticity we used an unbiased proteomic approach in combination with a statistical cell biology pipeline (the PRESTIGE) and show that transitions to mesenchymal-like forms following JAM3 depletion and/or TGFβ is coincident with upregulation of an EMP network comprised of canonical INS-AKT-FOXO, IKK-NFĸB, and Hippo-YAP/TAZ pathways. Critically, upregulation of the EMP network predicts mesenchymal states across cancers. The dynamics of this network suggest that EMP is coupled to a pro-inflammatory, insulin-resistant like state. Kinesin-1 is essential for upregulation of the EMP network, and inhibition of Kinesin-1 downregulates IKK-NFĸB activation and restores the signature of normal insulin signalling. By manipulating nuclear shape independently of Kinesin-1 we prove that changes in nuclear shape do not wholly explain the upregulation of the EMP network. Rather changes lead to the upregulation of sub-networks which ‘interlock’ with the EMP network. These sub-network encodes effectors of canonical INS-AKT, IKK-NFĸB, and Hippo-YAP/TAZ signalling pathways which interact with ‘hubs’ upregulated by Kinesin-1. Mechanistically, we propose changes in cell shape, such as loss of cell-cell adhesion in pre-mesenchymal cells alters MT organization and Kinesin-1 activity. Kinesin-1 activity and nuclear shape changes promote the activation of Ins-AKT-FOXO, Hippo-YAP/TAZ, and IKK-NFĸB pathways to drive EMP. Thus this system allows cells to sense the presence of disrupted tissue integrity, engage inflammatory responses, suppress growth, and promote mesenchymal morphogenesis. This pathway is essential in breast tissue to suppress tumorigenesis in response to environmental flux.

## Results

### Integrative omics demonstrates JAM3 is a tumour suppressor in breast cancer that regulates nuclear shape *in vivo*

Dysmorphic nuclei are a hallmark of cancer cells (Chow et al., 2012). Thus we identified genes whose levels correlated with changes in nuclear shape in 1,000 invasive human breast tumours (Curtis et al., 2012). We described the shape of single cancer cell nuclei, and the morphological heterogeneity of tumour cell nuclei *in vivo* by quantifying six shape features of tumour cell nuclei, but not the nuclei of lymphocytes or stromal cells, in haematoxylin and eosin (H&E) stained sections (Table S1; Figure 1A). For all features, a higher value indicates more irregular/elongated morphology, whilst a low value indicates a regular/round morphology. On average, there were 6 × 10^4^ cancer cells in the H&E sections in each tumour, and we used three statistics – median, standard deviation (sd), and skewness – to characterize the distribution of nuclear shape features in a tumour (Figure 1A). In total, 18 nuclear morphology scores were obtained for each tumour. Batch effects arising from different hospital sites and different staining and processing procedures were corrected for (Methods).

**Figure 1.**
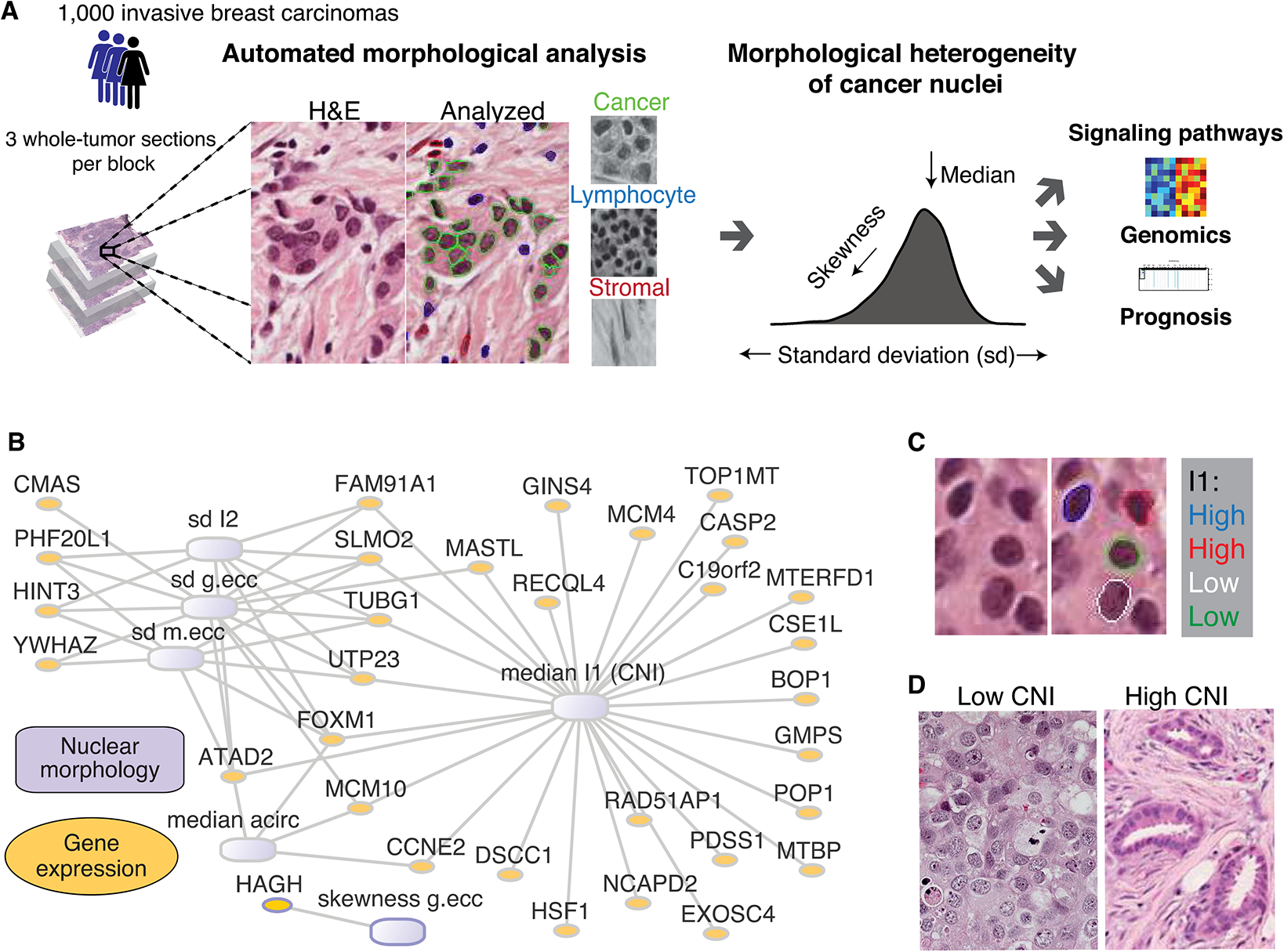
Computational pipeline for the integrative analysis of morphological and molecular data in human breast tumours. (A) Automated image analysis enables identification of three cell types and quantification of morphological features from an H&E whole tumour section. The distribution of each morphological feature for all cancer nuclei in a tumour image can be summarized using statistical descriptors: median, standard deviation, and skewness. These morphological scores can then represent tumour morphological profiles, which are then associated with genomic aberrations, gene expression data and patient prognosis. (B) Morphology-Expression hybrid Network representing significant associations between 18 cancer nuclear morphology scores and 1,000 cis-acting genes (feature name sd g.ecc: standard deviation of geometric eccentricity; median m.ecc: median value of moment eccentricity). (C) Illustrative examples of cancer nuclei with high and low value of image moment I1. (D) Illustrative examples of tumours with high and low CNI.

To identify cancer-driven molecular changes, we first identified the top 1,000 cis-driven genes based on association between DNA copy number alterations and mRNA expression across the 1,000 invasive breast cancers (Curtis et al., 2012). We then determined how the expression and copy number of these genes correlated with nuclear morphology and cell-to-cell variability in nuclear shape. The strongest gene expression-nuclear shape associations were used to construct a hybrid network (q<0.01 of false discovery rate (FDR); Spearman correlation corrected), with the core network module connecting 6 morphological scores and 38 gene expression probes representing 32 unique genes (q<0.005, Figure 1B). Nodes represent morphological scores or genes, and edges denote significant correlations. Given these genes were pre-selected based on their copy-number and gene expression profiles, this network effectively links DNA copy number, gene expression, and nuclear morphology.

Topology analysis of this network revealed that a morphological score “median I1” was positioned as the best-connected network hub (Methods). This feature describes rotational stability given the shape of an object. Intuitively, an object with asymmetric shape has a high I1 value. Cells and tumour tissues with different median I1 values are shown in Figures 1C and 1D. We henceforth refer to this morphological score as the Cancer Nuclear Instability index (CNI). Notably, “instability” here refers to a morphological property, as opposed to genomic instability. Well-differentiated epithelial-like cells have high CNI, suggesting CNI is prognostic (Figure 1D). Consistent with the idea that low CNI is a marker of cancer progression and aggressiveness, this score has a strong negative association with the mRNA levels of genes previously implicated in breast tumorigenesis as oncogenes including MCM4 (Choy et al., 2016), MCM10 (Baxley and Bielinsky, 2017), FOXM1 (Gartel, 2017), CCNE2 (Yasmeen et al., 2003), and MASTL (Marzec and Burgess, 2018) (Figure 1B). In addition, heterogeneity measures including standard deviation and skewness of morphology scores were also identified in this core network module; indicating that gene expression in single cells correlates with the extent of nuclear morphology heterogeneity in breast tumour cells *in vivo*. Thus both median changes in nuclear morphology as well as cell-to-cell variability in nuclear morphology are coupled to transcriptional upregulation of known oncogenes.

To identify putative tumour suppressors, we performed correlation analysis of CNI and genome-wide mRNA expression revealing 735 significant correlations (FDR correction q<0.01). CNI positively correlated with the mRNA expression of a number of genes encoding components of cell-cell adhesions, among which is JAM3 (Junctional Adhesion Molecule 3) whose mRNA was highly correlated with CNI (correlation=0.37, q-value 2.9 × 10^−6^) (Figure 2A; Table S2). JAM3 mRNA expression was strongly cis-driven (FDR corrected Spearman correlation among genome-wide associations between copy number and expression q-value=1 × 10^−7^), and was predominately lost in breast cancer (13% copy-number loss and only 0.8% gains; Figure 2B). The expression of JAM3 mRNA was lower in more aggressive Basal, HER2, and Luminal B subtypes compared to the less aggressive Luminal A subtype and “normal” subtypes (Figure 2C). JAM3 mRNA expression differed significantly in the 10 subtypes, among which “IntClust4” and “IntClust10” subtypes had the highest and lowest levels of JAM3 mRNA respectively (Figure 2D). IntClust4, which had high JAM3 mRNA expression, is the ‘CNA-devoid’ (no copy number alteration) group with favourable prognosis and a significant proportion of cases presented with extensive lymphocytic infiltration (Curtis et al., 2012). IntClust10, which had the lowest JAM3 mRNA expression, included the majority of Basal-like samples with high genomic instability. JAM3 mRNA expression was also lower in invasive ductal carcinoma (IDC) compared to less aggressive invasive lobular carcinoma (ILC) histology type (Figure 2F). The expression of JAM3 mRNA was also negatively correlated with tumour grade (Figure 2F). Thus, JAM3 expression scales with cancer aggressiveness and changes in nuclear shape.

**Figure 2.**
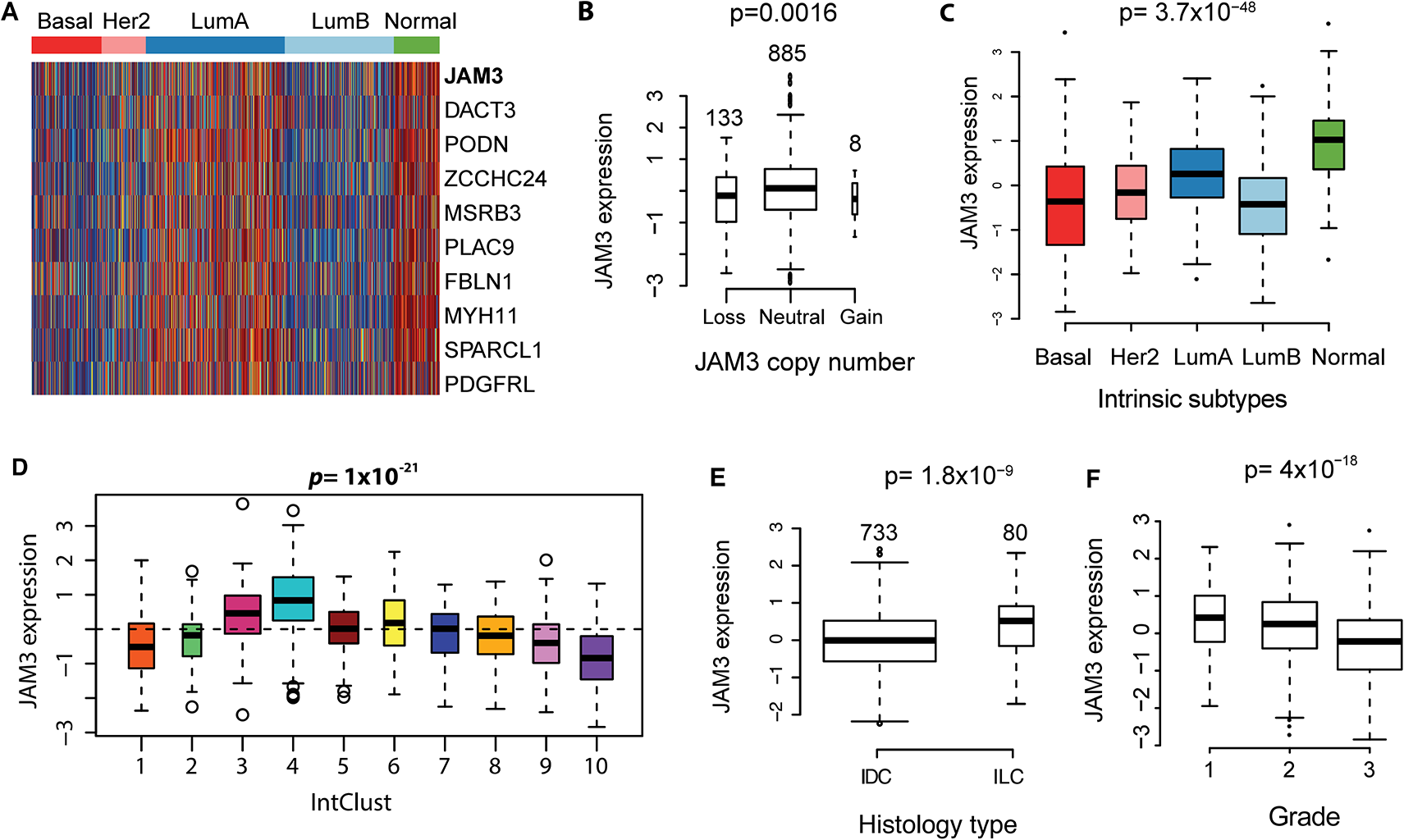
JAM3 as a potential driver of morphological heterogeneity. (A) Heatmap of RNA expression of genes with highest positive correlation with CNI, samples were annotated with intrinsic subtypes. (B) Boxplot showing the difference in JAM3 gene expression according to JAM3 DNA copy number status. (C) Boxplot showing the difference in JAM3 gene expression according to intrinsic subtypes. (D) Boxplot showing the difference in JAM3 gene expression according to the 10 integrative subtypes. (E) Boxplot showing the difference in JAM3 gene expression according to two main histology subtypes. IDC: Invasive Ductal Carcinoma; ILC: Invasive Lobular Carcinoma. (F) Boxplot showing the difference in JAM3 gene expression according to tumour grades.

### JAM3 regulates nuclear shape and microtubule organization in normal epithelial cells

We used quantitative single cell morphological analysis (Bakal et al., 2007) to assess how depletion of JAM3 affected the cell and nuclear shape of non-transformed MCF10A mammary epithelial cells. To this end, we used single or pooled small interfering RNAs (siRNAs) that effectively reduced JAM3 mRNA and/or protein. Two different pools (siGENOME or “siG”, and On-Target Plus or “OTP”) reduced JAM3 mRNA levels by 95 percent, and all individual siRNAs that constituted each pool reduced JAM3 mRNA levels between 75 and 97 percent (Figure S1A). Consistent with previous reports (Aurrand-Lions et al., 2001; Economopoulou et al., 2009; Lamagna et al., 2005; Zen et al., 2004), JAM3 protein is localized to the cell-cell junctions of MCF10A non-transformed epithelial cells, and the siG pool reduced JAM3 to virtually undetectable levels as judged by immunofluorescence (Figure 3A). The siG and OTP pools effectively depleted JAM3 protein, but not the related proteins JAM-A, as judged by quantitative mass spectrometry (Figure S1B) validating the on-target nature of these reagents. Although both siRNA pools and all individual siRNAs from each pool are highly effective at reducing JAM3 mRNA, for the majority of our experiments we used the siG pool and/or a single siRNA, OTP5, to deplete JAM3.

**Figure 3.**
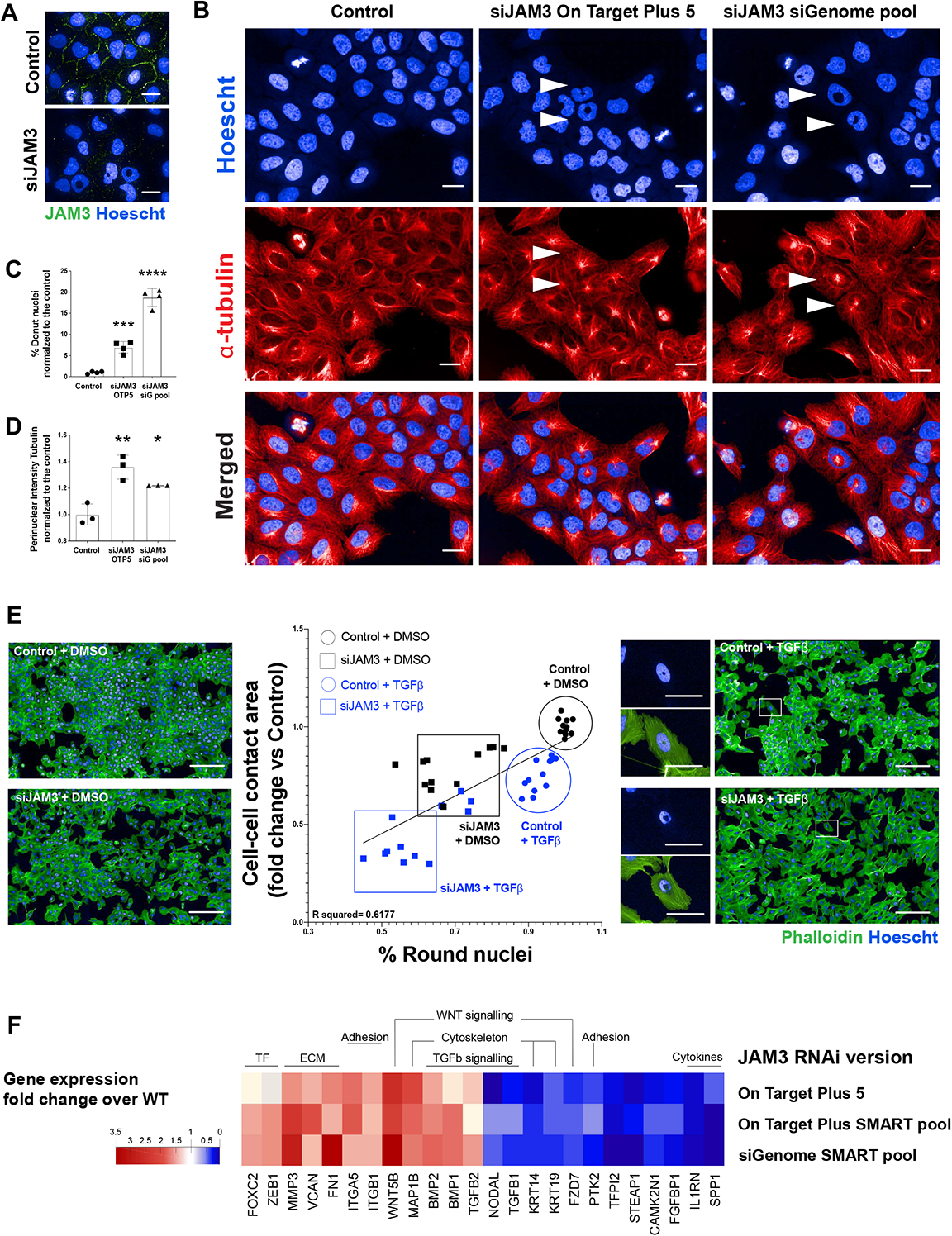
JAM-3 regulates nuclear and microtubule organization in epithelial cells. (A) Control or JAM3-depleted cells were labelled with Hoechst (blue) and immunostained for JAM-3 (green). Scale bars, 20um. (B) Representative images of the ‘donut’ nuclei (blue; labelled with Hoescht) and the microtubules (red; immunostained with α-tubulin) characteristic of MCF10A cells transfected with 2 different siRNAs targeting JAM3. Scale bars, 50um. (C, D) Quantification of perinuclear α-tubulin intensity (C) and percentage of cells with nuclear donuts (D) in control and JAM-3 depleted cells. Graph shows the representative data of 3 different experiments (n > 12.000 cells). Comparisons among groups were performed by one-way ANOVA (Newman-Keuls multiple comparison test; ∗p < 0.05, ∗∗p < 0.01, ∗∗∗p < 0.001, ∗∗∗∗p < 0.0001). (E) Average neighbour fraction (cell-cell contact area) were positively correlated with the percentage of cells with round nuclei in wild-type and JAM3 depleted cells, non treated or treated with TGFb for 2 days. Graph shows the result of 3 independent experiments where each point is the well average of at least 700 cells. Images are showing examples of each treatment labelled with Phalloidin (green) and Hoescht (blue) and cropped sections are showing the nuclei deformation after TGFb treatment of wild-type cells or of JAM3 deleted cells. Scale bars, 200um. (F) Heat map showing relative gene expression panel after treating MCF10A cells with 3 different siRNAs targeting JAM3.

JAM3 depletion in MCF10A cells resulted in striking changes in nuclear morphology. The vast majority of nuclei of normal MCF10A cells were shaped like smooth ovals (Figure 3B). In contrast, the nuclei of JAM3 depleted cells had highly irregular, banana-like shapes, and a proportion had large holes in the nuclei – appearing toroidal (Figure 3B). We termed the nuclei with large holes “donut” nuclei. Using a linear classifier, we automatically identified donut nuclei in cellular populations (Methods). Normal cell populations had ∼1% of nuclei that could be classified as donuts, but JAM3 depletion increased the proportion of donut nuclei population to ∼10-20% (Figure 3C). Inhibition of JAM3 by 3/4 individual siRNAs from the siG pool, or 2/4 siRNAs from the OTP pool (Figure S1C), led to significant increases in donut nuclei. Because these individual siRNAs target different sequences this. The phenotypic similarity of cells treated with different siRNAs targeting JAM3 that effectively deplete JAM3 mRNA and protein demonstrates the specificity of JAM3 knockdown, and eliminates the possibility this phenotype can be attributed to off-target effects.

Depletion of JAM3 by siRNA also resulted in the dramatic reorganization of MTs. Whereas the MTs of wild-type cells were organized into parallel filaments that span the long axis of the cell and loosely grouped in the perinuclear region (Figure 3B), the MTs of JAM3 depleted cells were consistently organized into radial arrays with highly focused microtubule organizing centres (MTOC) (Figure 3B chevrons). JAM3 depleted cells had significant higher perinuclear MT intensity as judged my quantitative single cell analysis (Figure 3D). These data suggest that JAM3 depletion affects MT organization.

We observed the MTOC of JAM3 depleted cells was almost always positioned in the hole of nuclear donuts, or at the deepest point of invagination in banana nuclei (Figure 3B). Knockdown of JAM3 in other normal epithelial cell types, such as Retinal Pigmented Epithelial (RPE-1) cells, had identical effects on nuclear morphology (Figures S1D, S1E). Notably, depletion of JAM3 had little effect on the organization of the actin cytoskeleton, especially near the nucleus (Figure S1F, S1G).

### JAM3 depletion synergizes with TGFβ to promote Epithelial-Mesenchymal Plasticity

The change in MT organization, from partially aligned MT fibers characteristic of more epithelial cells to radially organized MTs with a strong MTOC that are similar to MTs of mesenchymal/fibroblast cells (Keating and Borisy, 1999) led us to investigate whether loss of JAM3 induced either a full or partial transition from an epithelial to a more mesenchymal state. To quantify the extent of this transition following JAM3 depletion we measured the cell-cell “contact area”, which describes the extent of cell-cell contact on a per cell basis, and cells in epithelial monolayers that are completely surrounded by other cells will have an NF of 1. As a control we treated both control and JAM3 depleted MCF10A cells with the canonical TGFβ inducer for 48 hrs (Miettinen et al., 1994). Both TGFβ or JAM3 depletion by themselves or together resulted in significantly decreased NF and disrupted colony formation, supporting the idea that JAM3 is necessary for maintenance of an epithelial state (∼75% of control; Figure 3E). NF and the percent of round nuclei in the population was correlated, as decreases in NF led to decreases in cells with spherical nuclei (Figure 3E). Notably, combined TGFβ stimulation and JAM3 depletion synergistically pushed epithelial-like MCF10A cells towards mesenchymal forms beyond either treatment alone (Figure 3E). Thus we conclude that JAM3 depletion promotes EMP and differentiation towards a mesenchymal state. Changes in MT and nuclear organization (MTNO) occur at the same time as mesenchymal differentiation induced by JAM3 depletion and/or TGFβ treatment.

The mesenchymal-like phenotype following JAM3 loss was confirmed by the observation that JAM3 deficient cells upregulate pro-mesenchymal mRNAs including Fibronectin-1 (FN), ITGA5, MMP3, BMP1, and MAP1B, and downregulate pro-epithelial mRNAs such as KRT14 and KRT19 (Figure 3F). We confirmed upregulation of individual mRNAs by qPCR (Figure S1H; Table S3).

### JAM3 is frequently lost in mesenchymal cancers

To understand if loss of JAM3 protein is a common occurrence during transitions to mesenchymal forms in cancer we examined the total protein abundances of 12,197 unique proteins across 375 unique cancer cell lines from the Cancer Cell Line Encyclopaedia (CCLE) (Ghandi et al., 2019). Of these, 324 had detectable levels of the epithelial marker CDH1, the mesenchymal marker VIM, and JAM3. 170/324 lines had high Vimentin/VIM and low CDH1 expression and could broadly be considered mesenchymal (Nusinow et al., 2020). 99/170 (58%) of these mesenchymal lines also had low JAM3 expression demonstrating that loss of JAM3 and increased VIM expression are often coincident during EMP in cancer cells (Figure S2A, Table S4). These data suggest that JAM3 and CDH1 loss are not mutually exclusive and function independently

To validate the notion that JAM3 depletion leads to EMP more directly, we tested whether JAM3 depletion resulted in non-invasive MCF10A cells gaining the ability to invade 3D collagen matrices where nuclei are compressed as they move through pores in the matrix; a characteristic of mesenchymal cells (Figure S2B, S2C). Whereas normal MCF10A cells do not invade collagen matrices, JAM3 depleted cells invaded the gels to heights of 60 μm in 24 hrs (Figure S2B S2C). Collagen invasion was dependent on Matrix Metalloproteinases (MMPs; Figure S2D, S2E).

### JAM3 is required for the morphogenesis of epithelial tissues

To assess the role of JAM3 in promoting epithelial morphogenesis, we investigated the ability of JAM3-deficient MCF10A cells to form organoid (acini) structures in solid matrices in mixtures of Matrigel and Collagen-I (Col-I; Methods). After four days in 3D matrices, normal MCF10A cells formed round organoids with a well-organized thin outer layer of Laminin-5 (Figure S3A, S3B). In contrast, JAM3 depletion (JAM3 OTP or siG) decreased both the number of cells in each organoid structure and the acini area (Figure S3A, S3B). JAM3 deficient organoids also formed small protrusive actin-rich ‘neurites’ which were basically non-existent in control cells (Figure S3B). The laminin-5 basement membrane in JAM3-depleted organoids was absent compared to wild-type, but FN was organised into large bundles that encapsulated the organoids (Figure S3A, S3B). We observed significant increases in FN intensity with respect to actin staining (Figure S3B). Over eight days, JAM3-depleted organoids developed into irregularly shaped bodies which often fused into larger multi-acinar structures (Figure S3C, S3D). These results demonstrate JAM3 is essential for mammary organoid morphogenesis in 3D.

### JAM3 suppresses the activity of YAP and TAZ

Both YAP and TAZ are structurally related proteins, with only partially overlapping functions, that translocate from the cytoplasm to nucleus in response to forces on the nucleus (Elosegui-Artola et al., 2017; Piccolo et al., 2014; Shiu et al., 2018), and regulate the transcription of genes that results in changes in cell shape, promote migration, proliferation, and invasion (Lamar et al., 2012; Overholtzer et al., 2006; Shao et al., 2014). We hypothesized that because JAM3 depletion alters nuclear morphogenesis, which coincides with differentiation into a mesenchymal-like state, this may coincide with activation of the YAP and/or TAZ transcriptional co-activators. In wild-type MCF10A cells cultured at medium to high densities, both YAP and TAZ are localized to the both the cytoplasm and the nucleus as determined by immunofluorescence staining (Figure S4A). However, the ratio of nuclear:cytoplasmic YAP and TAZ increased significantly after downregulation of JAM3 by RNAi (Figure 4A-C).

**Figure 4.**
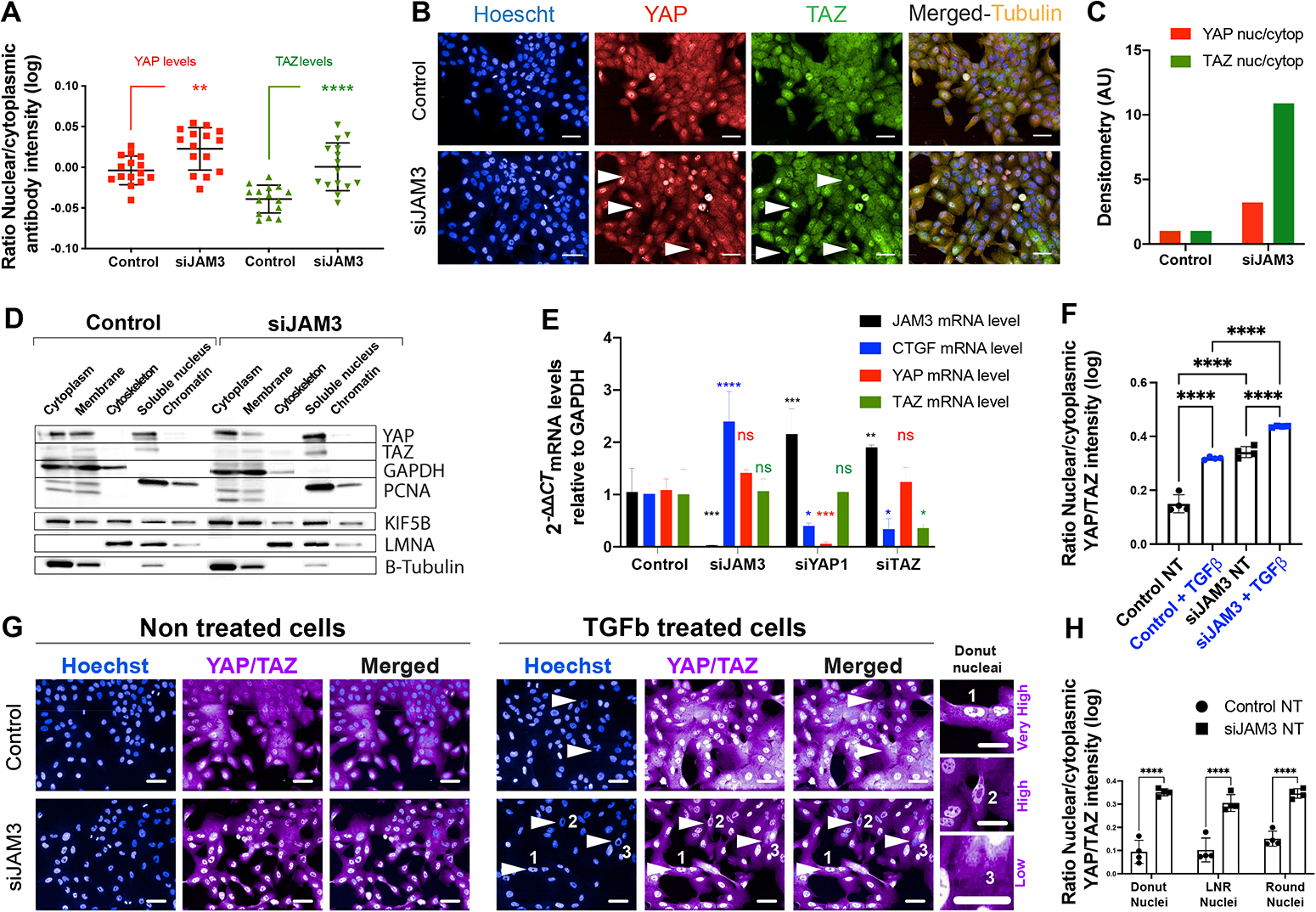
JAM3 regulates translocation of YAP and TAZ. (A) Quantification of the nuclear:cytoplasmic YAP or TAZ ratios for wild-type and JAM3-depleted cells (n > 15.000 cells). Data shows the results of 3 different experiments. Comparisons among groups were performed by one-way ANOVA (Newman-Keuls multiple comparison test) with Prism6 software (∗p < 0.05, ∗∗p < 0.01, ∗∗∗p < 0.001, ∗∗∗∗p < 0.0001). (B) Wild-type and JAM3-depleted cells were labelled with Hoechst (blue) and inmunostained for TAZ (green), YAP (red) and α-tubulin (yellow). Scale bars, 50um. (C) YAP and TAZ nuclear:cytoplasmic ratio as determined by biochemical fractionation in control and siJAM3 depleted (siG RNAi) cells. Results are from one experiment where 1.000.000 cells were lysed per condition. (D) Representative western blots of subcellular fractions for Wild-type and JAM3 depleted cells that were blotted for YAP and TAZ and for several loading controls: KIF5B (cytoskeleton fraction), PCNA and LMNA (soluble nuclear and Chromatin bound fractions) and GAPDH and β Tubulin (cytoplasmic and membrane fractions). (E) Relative mRNA levels of CTGF, JAM3, YAP and TAZ normalized to GAPDH for wild-type and JAM3, YAP, or TAZ depleted cells. Mean ± SD (n = 3 replicates/condition). (F) Quantification of the nuclear:cytoplasmic YAP/TAZ ratios for wild-type and JAM3-depleted cells non treated or treated with TGFb for 2 days. Graph shows the representative data of 2 different experiments (n > 3.000 cells per condition). Comparisons among groups were performed by two-way ANOVA (Šídák’s multiple comparisons test) with Prism6 software (∗p < 0.05, ∗∗p < 0.01, ∗∗∗p < 0.001, ∗∗∗∗p < 0.0001). (G) Wild type cells or JAM3 depleted cells non treated or treated with TGFb were labelled with cells were labelled with Hoechst (blue) and inmunostained for YAP/TAZ (purple). Arrows and numbers are showing 3 different donut nuclei with 3 different levels of YAP/TAZ translocation. Scale bars, 50um. (H) Non treated control and JAM3 depleted cells were classified according to their nuclei shape and YAP/TAZ translocation was measured. Graph shows the representative data of 2 different experiments (n > 3.000 cells per condition). Comparisons among groups were performed by two-way ANOVA (Tukey’s multiple comparisons test) with Prism6 software (∗p < 0.05, ∗∗p < 0.01, ∗∗∗p < 0.001, ∗∗∗∗p < 0.0001).

To confirm that JAM3 depletion results in a change in the subcellular localization of YAP and TAZ, we performed biochemical fractionation of mock-treated and JAM3 depleted cells, and quantified the levels of YAP and TAZ in each fraction by Western blotting. Indeed, JAM3 depletion resulted in a significant accumulation of both YAP and TAZ protein in the nuclear fraction (Figure 4C, D), and depletion of YAP and TAZ from membrane fractions (Figure 4D). JAM3 depletion also resulted in YAP and TAZ activation as well as increased levels of CTGF mRNA (Figure 4E), a canonical target of YAP/TAZ mediated transcription (Zhao et al., 2008). Stimulation by TGFβ also upregulated YAP/TAZ nuclear translocation and did so in a manner synergistic with JAM3 depletion (Figure 4F, 4G). Thus YAP/TAZ is activated during upregulation of EMP by JAM3 depletion and TGFβ stimulation.

We next sought to determine if increased YAP and/or TAZ nuclear translocation following JAM3 depletion are responsible for altered organoid morphogenesis during EMP, such as the formation of invasive protrusions. We thus knocked down YAP or TAZ in control or JAM3-deficient MCF10A cells cultured in 2D or as organoids. Both YAP and TAZ siRNAs were highly effective and specific at knocking down their targets (Figure S4A, S4B, S4C). But while co-depletion of JAM3 and YAP1 resulted in organoids with multiple actin-rich protrusive structures which resemble organoids depleted of JAM3 alone (Figure S4D, SE), co-depletion of TAZ with JAM3 completely abrogated the formation of actin-rich invasive structures as judge by neurite number and total neurite length (Figure S4D, SE). In contrast, depletion of YAP prevented remodelling of FN surrounding the acini (Figure S4E). Thus TAZ, but not YAP, is essential for the generation of protrusive structures made by abnormal organoids following JAM3 depletion. But YAP is required for ECM remodelling following JAM3 depletion.

### YAP/TAZ activation is not due to changes in nuclear shape

Changes in nuclear morphology have been linked to YAP/TAZ activation. To assess whether changes in nuclear morphology following JAM3 depletion and/or TGFβ stimulation were causal to YAP/TAZ translocation, we examined YAP/TAZ nuclear levels in wild-type, and JAM3 depleted, cells that were classified into different nuclear shape classes (‘round, low-nuclear roundness, and donut). All nuclear shape classes had equivalent levels of increased YAP/TAZ translocation (Figure 4H). Thus we conclude that increased YAP/TAZ translocation is not due to changes in nuclear shape during EMT.

### SYNE4 and KIF5B depletion rescue the phenotypes of JAM3 depletion

To determine if cytoskeletal reorganization was responsible for nuclear deformation in JAM3 deficient cells we systematically depleted all known components of the SUN-KASH complexes, which bridge the cytoskeleton and the nuclear lamina (Starr and Fridolfsson, 2010), with siRNA pools in JAM3-depleted. Because SYNE4 and KIF5B, the heavy chain of the MT-bound motor Kinesin-1, physically interact to regulate nuclear position and chromatin mobility, we also depleted SYNE4 and KIF5B in combination with JAM3. Only co-depletion of JAM3, SYNE4, and KIF5B (hereafter referred to as the “triple” knockdown) led to a complete reversion of the abnormal nuclear shape (Figure 5A) and MT reorganization (Figure 5B) observed in JAM3 deficient cells alone. Specifically, donut nuclei were clearly observed in all conditions except the triple knockdown (Figure 5C, chevrons. Triple knockdown in green box). Co-depletion of JAM3 with SYNE1, SYNE2, or SYNE3 did not rescue the phenotypes of JAM3 depletion (Figure 5A, 5B) suggesting that actin or intermediate filaments do not play a role in nuclear deformation in JAM3 depleted cells. These results support the idea that disrupting the complexes that bridge the MTs and the nucleus by depleting SYNE4 and KIF5B can prevent MTs from deforming the nucleus in JAM3 depleted cells.

**Figure 5.**
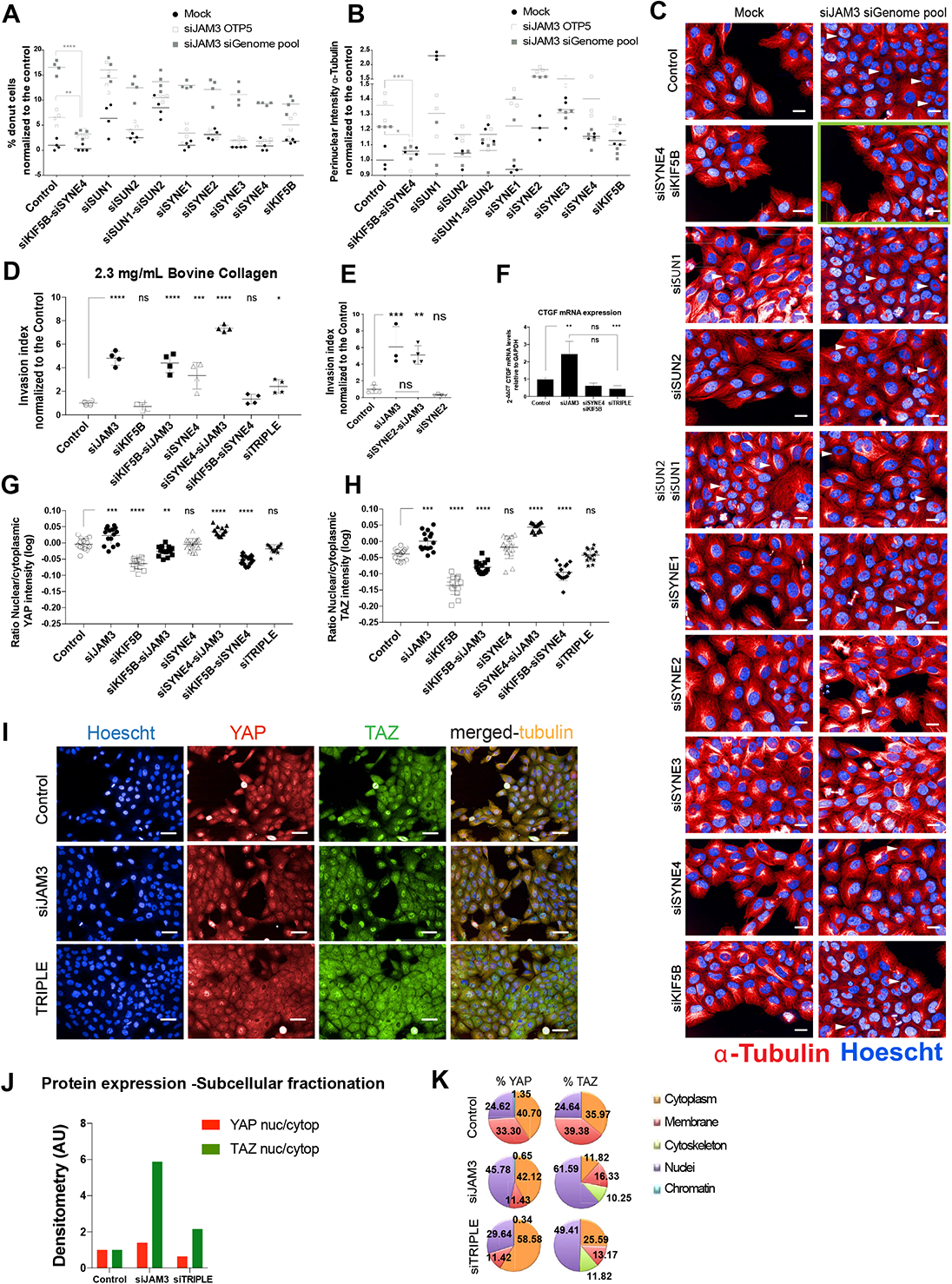
Depletion of KIF5B and SYNE4 prevent EMP. (A) Percentage of cells with nuclear donuts and (B) quantification of Intensity of perinuclear α-tubulin in cells treated with JAM3 siRNA alone, or in combination with a panel of siRNAs targeting almost all the LINC components (n > 4.000 cells). Data shows the results of one experiment. Comparisons among groups were performed by one-way ANOVA (Newman-Keuls multiple comparison test) with Prism6 software (∗p < 0.05, ∗∗p < 0.01, ∗∗∗p < 0.001, ∗∗∗∗p < 0.0001). (C) Representative images of the nuclei (blue, Hoechst) and the MTOC (red, α-tubulin) of MCF10A cells transfected with a panel of siRNAs targeting several LINC components alone or in combination with JAM3. Scale bars, 50um. (D) Invasion index of MCF10A cells depleted of single genes, or genes in combination. Graphs show the representative data of 2 different experiments (n > 5.000 cells). Comparisons among groups were performed by one-way ANOVA (Newman-Keuls multiple comparison test) with Prism6 software (∗p < 0.05, ∗∗p < 0.01, ∗∗∗p < 0.001, ∗∗∗∗p < 0.0001). (E) Invasion index of MCF10A cells depleted of single JAM3 and/or SYNE2 into a Collagen matrix. Graph shows the data of one experiment (n > 5.000 cells). Comparisons among groups were performed by one-way ANOVA (Sidak’s multiple comparison test) with Prism6 software (∗p < 0.05, ∗∗p < 0.01, ∗∗∗p < 0.001, ∗∗∗∗p < 0.0001). (F) Relative mRNA levels of CTGF normalized to GAPDH for wild-type, JAM3 depleted, KIF5B+SYNE4 depleted, and Triple knockdown cells. Mean ± SD (n = 3 replicates/condition). (∗p < 0.05, ∗∗p < 0.01). (G-I) Wild-type, JAM3-depleted cells single or in combination with KIF5B and/or SYNE4 RNAi were labelled with Hoechst (blue) and inmunostained for TAZ (green), YAP (red) and α-tubulin (yellow). (G) Quantification of the nuclear/cytoplasmic YAP ratio after the RNAi treatments. (H) Quantification of the nuclear/cytoplasmic TAZ ratio after the RNAi treatments. (n>15.000 cells). Data shows the results of 3 different experiments. Comparisons among groups were performed by one-way ANOVA (Newman-Keuls multiple comparison test) with Prism6 software (∗p < 0.05, ∗∗p < 0.01, ∗∗∗p < 0.001). (I) Representative images of the wild-type, JAM3 and Triple depleted cells. Scale bars, 50um. (J) YAP and TAZ nuclear:cytoplasmic ratio as determined by biochemical fractionation in wild-type, JAM3 and Triple depleted cells. (K) YAP and TAZ subcellular localization as determined by biochemical fractionation in control, JAM3 depleted, and triple knockdown cells. Pie charts represent the percentage of YAP and TAZ protein localization in each fraction determined by WB of the different extracts for each RNAi treatment. Results are from one experiment where 1.000.000 cells were lysed per condition.

To determine if the rescue of nuclear shape phenotypes in JAM3 depleted cells following depletion of SYNE4 and KIF5B also reversed the engagement of pro-invasive programmes, we measured 3D invasion in: 1) normal cells; 2) cells depleted of JAM3, KIF5B, or SYNE-4 alone; 3) cells co-depleted of JAM3 and KIF5B or SYNE-4; and 4) cells co-depleted of JAM3, KIF5B, and SYNE-4 (triple knockdown). Whereas depletion of JAM3 alone, or co-depletion of JAM3 and KIF5B or SYNE-4, led to increased invasion, the triple knockdown cells did not invade (Figure 5D). Notably, co-depletion of the actin binding protein SYNE-2 and JAM3 did not prevent invasion (Figure 5E). Thus, the function of both KIF5B and SYNE-4 are required for mesenchymal phenotypes, and in particular the increased invasiveness, associated with depletion of JAM3.

In JAM3 depleted cells, KIF5B and SYNE4 are also required for elevated CTGF mRNA levels (Figure 5F), and nuclear translocation of YAP (Figure 5G, I) and YAZ (Figure 5H, I). We confirmed that SYNE4 and KIF5B promote increased YAP and TAZ nuclear translocation by subcellular fractionation (Figure 5J).

### Inhibition of Kinesin-1 inhibits mesenchymal morphogenesis

To validate the role of Kinesin-1 function in promoting EMP such as following JAM3 knockdown and eliminate any possibility that KIF1B siRNA had off-target effects we treated JAM3, or JAM3/SYNE4 deficient cells, with Rose Bengal Lactone (RBL). RBL is a fluorescein compound that binds Kinesin-1 at the L8/β5 pocket and inhibits the binding of Kinesin-1 to MTs (Hopkins et al., 2000). We also used RBL to inhibit Kinesin-1 activity in control TGFβ treated cells, or JAM3 depleted and TGFβ treated cells. We determined the extent of EMP in each population by quantifying “cell-cell contact area”. Both JAM3 and TGFβ treated cell have ∼70% of the contact area compared with normal cells (Figure 6A), due to disruption of colony-like structures (Figure 6B). JAM3 depleted cells treated with TGFβ have 30% of cell-cell contacts compared to control, indicative of an almost complete dispersal of colonies and full-blown EMT. Consistent with our previous observations, cell-cell contact in cells co-depleted of SYNE4 and KIF5B with JAM3 (siTriple) was similar to wild-type. Chemical inhibition of Kinesin-1 by RBL also restored cell-cell contact in JAM3 depleted and/or TGFβ treated cells to levels of wild-type. Thus we conclude Kinesin-1 activity is required for mesenchymal morphogenesis in JAM3 depleted and/or TGFβ treated cells.

**Figure 6.**
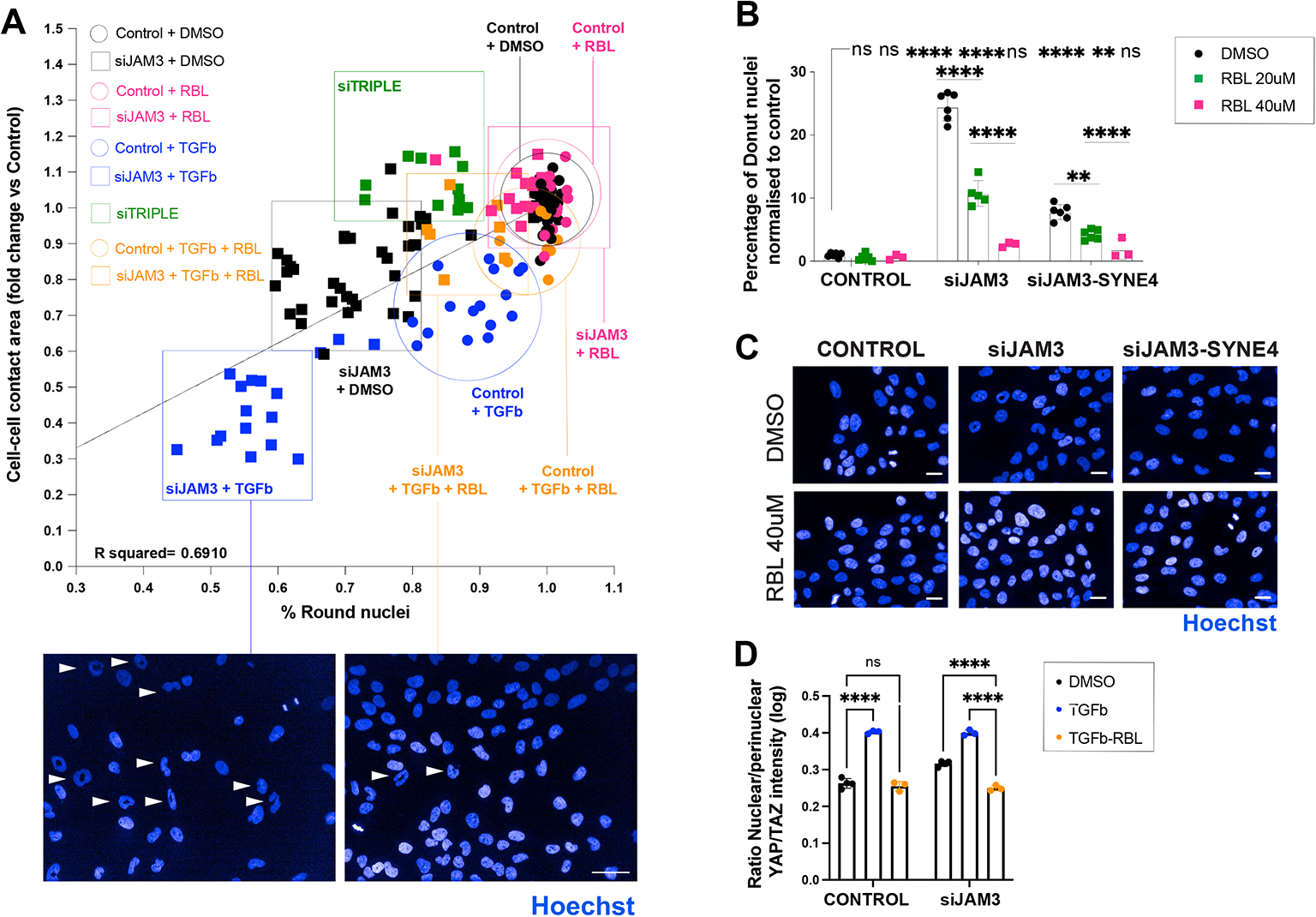
Chemical inhibition of Kinesin-1 prevents EMP. (A) Average cell-cell contact area were positively correlated with the percentage of cells with round nuclei across a panel of different RNAi and/or chemical treatments. Data shows the results of at least 3 different experiments where each point is the well average of at least n > 700 cells. Pics are showing the decrease of round nuclei population after JAM3 depletion and TGFb treatment and the rescue of this deformability and neighbour fraction by RBL at 40uM for 2 days. R squared was determined by using Prism6 software (simple linear regression). (B) Quantification of YAP/TAZ nuclear translocation levels in Wild-type or JAM3-depleted cells non treated or TGFb treated or TGFb + RBL treated. Graph shows the representative data of 2 different experiments (n > 3.000 cells per condition). Comparisons among groups were performed by two-way ANOVA (Šídák’s multiple comparisons test) with Prism6 software (∗p < 0.05, ∗∗p < 0.01, ∗∗∗p < 0.001, ∗∗∗∗p < 0.0001). (C,D) MCF10A cells mock transfected, transfected with JAM3 RNAi single or in combination with SYNE4 RNAi and treated with DMSO or RBL at the indicated concentrations for 48 hours. (C) Percentage of cells with nuclear donuts (n > 3.000 cells) where RBL mirrors the effects of KIF5B RNAi in the siJAM3-siSYNE4-RBL condition. Comparisons among groups were performed by one-way ANOVA (Newman-Keuls multiple comparison test) with Prism6 software (∗p < 0.05, ∗∗p < 0.01, ∗∗∗p < 0.001, ∗∗∗∗p < 0.0001). (D) Representative images of the nuclei (blue) of all the conditions. Scale bars, 50um.

RBL rescued nuclear morphology defects in both JAM3 and JAM3/SYNE4 depleted cells (Figure 6), which correlated with the reversion of mesenchymal state to an epithelial state (as judged by “contact Area with neighbours” feature, Figure 6). These data demonstrate that Kinesin-1 activity is essential for both EMP and nuclear shape changes that underpin EMP in JAM3 or TGFβ-treated cells.

### JAM3 depletion affects LMNA levels and phosphorylation

The defects in nuclear morphogenesis following JAM3 depletion were highly reminiscent of nuclear morphology defects observed in cells with dysfunctional Lamin A (LMNA), such as progeric cells (Eriksson et al., 2003). Antibodies specific to LMNA and LMNB were used to detect either the total levels of nuclear LMNA and LMNB (Figure S5A). The ratio of LMNA or LMNB localized to the nuclear ring region compared to the inner ring region was measured as a proxy for the amount of each protein that is incorporated into the nuclear lamina (Figure S5B, S5C). Depletion of LMNA, but not LMNB, increased the percentage of misshapen nuclei in the population, and that of donuts in particular (Figure S5D, SE). But while combined depletion of LMNA and JAM3 also led to a marked increase in the percentage of misshapen nuclei in the population, the nuclei of JAM3 and LMNB deficient cells were the same shape as wild-type nuclei (Figure S5E chevrons). Thus the effects on JAM3 depletion on nuclear shape are at least in part due to decreased level of LMNA.

Depletion of JAM3 affected both the outer:inner nuclear ratio of LMNA and LMNB (Figure S5F, S5G). Combined depletion of SYNE4 and KIF5B in the background of JAM3 depletion (the “triple” knockdown) restored the levels of LMNA but not LMNB (Figure S5G) in the nuclear lamina. Interestingly, similar to SYNE4/KIF5B depletion, depletion of YAP/TAZ also rescued the decreased LMNA, but not LMNB levels in JAM3 deficient cells (Figure S5G). We conclude lowered LMNA levels during mesenchymal morphogenesis may in part be due to YAP/TAZ transcriptional activity. Thus nuclear shape changes may be a consequence, but not cause, of YAP/TAZ activation.

Because phosphorylation of LMNA at serine 22 (hereafter referred to a pLMNA) promotes the dissociation of LMNA filaments (Buxboim et al., 2014; Heald and McKeon, 1990), we probed Ser22 phosphorylation following JAM3, SYNE4 and/or KIF5B. Depletion of JAM3 significantly increased the total levels of pLMNA (Figure S5H, S5I). Increased pLMNA levels were completely rescued in the triple knockdown. In summary, these data suggest that JAM3 regulates the LMNA phosphorylation status, stability, and incorporation into the nuclear lamina.

### A statistical cell biology framework, PRESTIGE, identifies an EMP network

To understand the basis for how changes in cell shape and MTNO may regulate epithelial-mesenchymal plasticity (EMP) we global quantitative proteomes (Roumeliotis et al., 2017) (Table S5). We compared the total protein intensities of JAM3 depleted cells (Mesenchymal), TGFβ treated cells (Mesenchymal), or TGFβ treated JAM3 depleted cells (Mesenchymal), among each other and also against the intensities from non-treated wild-type cells (Epithelial) To identify pathways which are both commonly and uniquely regulated in each condition, we developed PRESTIGE (Pathway RESToration via Integrated imaGe-Expression datasets) a statistical cell biology framework which combines quantitative imaging data, peptide levels, and protein-protein interaction (PPI) datasets (Methods). Briefly, PRESTIGE functions by grouping different experiments into classes based on imaging data and cellular phenotypes (versus experimental labels alone), and we identify peptides whose levels are significantly up/down regulated in each individual class.

Firstly, 2-way ANOVA was applied to quantify the variances across phenotypes and identified 260 proteins as upregulated upon JAM3 depletion and/or TGFβ treatment. Of these, 223 (85%) satisfied our criteria (Methods) for inclusion into a list of proteins consistent upregulated in these conditions (i.e. versus being strongly upregulated in one condition, but unaffected or even downregulated significantly in others). These correspond to 1.08% of all proteins in the dataset.

Using PRESTIGE we generated networks of proteins upregulated during EMP (Figure S6, Table S5). Following JAM3 depletion, with or without TGFβ treated, we noted elevated levels of proteins that comprised members of Insulin signalling (e.g. INS, IRS1, IRS2, PIK3CD, PIK3R2, AKT3, PTEN, and FOXO1) and canonical NFĸB (e.g. RELA, NFĸBIA, and IKBKB) pathways. Indeed, proteins upregulated during EMP are significantly enriched in the KEGG_’Insulin_Resistance’ category which contains both canonical Insulin-AKT and NFKB signalling proteins (HSA04931;p-value of 0.01121, q-value of 0.09754) (Table S7).

Insulin itself had 17 PPIs and was the most connected protein in the EMP network of upregulated proteins, while PTEN, FOXO1, IRS1, IKBKB, and AKT3 all had at least 10 PPIs (Methods). Because MCF10A cells are incapable of insulin production, and it is added to the culture medium as a supplementary growth factor, we hypothesized that elevated insulin levels are due to changes in insulin uptake, but not production.

We plotted the 48 proteins in the PPI and functional neighbourhood for JAM3 and INS that are upregulated in TGFβ and/or JAM3 depleted cells as a network where nodes were assigned to a subcellular locations (Figure 7A). The nodes had the same colour pattern as in Fig S6, with 25 members of KEGG Insulin_Resistance (HSA04931) highlighted using red boundaries around the nodes. We term this a ‘core EMP’ network.

**Figure 7.**
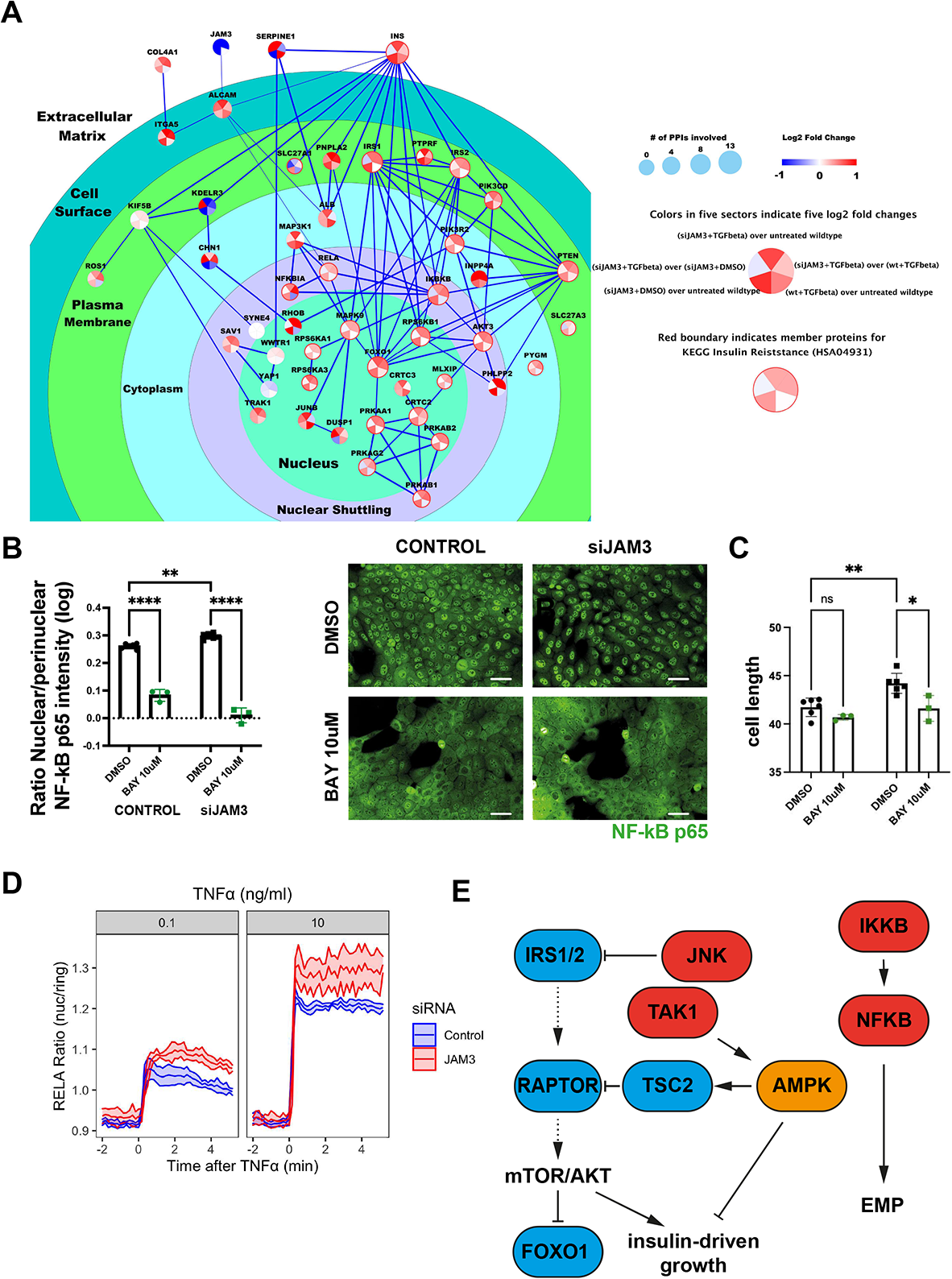
The core EMP network drives inflammation. (A) The core EMP network. Each protein is represented by one circle node evenly split into five sectors. The colour of the five sectors is proportional to the log2 transformation of fold changes from five different comparisons indicated in the legend section. All nodes showed significant up-regulation (red color as indicated in the legend) in at least one of the five comparisons, and more than half were up in at least three comparisons. Specifically, 25 of the 48 proteins belonged to KEGG Insulin Resistance pathway (HSA04931) and they were highlighted by red boundary around the node. One edge summarizes all PPIs between the two connected nodes, as recorded in the STRING database. The width of each edge is proportional to the overall confidence score given by STRING. The size of each node is proportional to the number of edges connected to it. The primary subcellular locations for the proteins were obtained from InnateDB Ver.5.4, where cell surface was defined by Gene Ontology term GO:0009986 and differentiated from plasma membrane. The localization layer of nuclear shuttling was defined by domain expert. The localizations were visualized using rings and circles with different colours. (B) Wild-type and JAM3-depleted cells were treated with DMSO or the IKKi BAY 11-7082 for 75min and then stimulated with 10ng/mL for 30min, fixed and inmunostained for NFκB p65. Graph shows the representative data of 2 different experiments (n > 8.000 cells per condition). Comparisons among groups were performed by two-way ANOVA (Turkey’s multiple comparison test) with Prism6 software (∗p < 0.05, ∗∗p < 0.01, ∗∗∗p < 0.001, ∗∗∗∗p < 0.0001). On the right are representative images showing NFκB p65 (green) in TNFα treated cells. Scale bars, 50um. (C) IKK is partially responsible for cell shape changes during EMP. JAM3 depletion causes increases in cell-length which are blocked by IKK inhibition (C) RELA ratio (nuc/ring region RELA-GFP intensity) in RPE1 cells expressing AAV mRuby-PCNA with CRISPR/CAS9 RELA-GFP. Per condition, ribbons show the 95% confidence interval with a central line showing the mean RELA ratio across six technical replicates. Cells were transfected 48 hr prior with non-targeting or JAM3 siRNA then imaged for 2 hr pre-TNF treatment and 5 hours post-TNF (0, 0.1 or 10 ng/ml). (D) Cartoon representation of signalling networks activated during EMP. JNK activity inhibits IRS1. TAK1 activates AMPK, which can in turn activates TSC2 that inhibits mTOR and AKT. IKK activates p65/RELA which is partially responsible for EMP.

Based on the concomitant upregulation of NFKB modulators and components of the insulin pathway − we hypothesized that activation of pro-inflammatory signalling is likely downregulating AKT signalling during EMP. To test this hypothesis, we examined the phosphoproteomes of control (epithelial-like) and JAM3 depleted and/or TGFβ treated (mesenchymal-like) cells (Table S8). As expected, during EMP there was significant alteration in the phosphorylation of proteins in Tight Junction (q=0.000468), TGFβ (q=5.47E-06), and cell cycle (q=1.47E-08) KEGG or Wikipathways categories (Table S9).

In mesenchymal cells, we observed extensive phosphorylation on MAPKs which activate both JNK and NFĸB signalling during EMP including NFĸB activators IKBKG, MEKK3/MAP3K3; JNK activators MAP2K4/JNKK1, MAP3K4/MEKK4, MAP3K6/AKK2; and dual JNK/NFKB activators MAP3K7/TAK1, MAP3K1/MEKK1, and MAP3K8/COT (Table S8). These data demonstrate EMP occurs concomitantly with an increased in inflammatory signalling – in line with our previous studies in breast cancer cells (Sero et al., 2015).

We confirmed the upregulation of IKK-NFĸB signalling following JAM3 depletion in two different models. JAM3 depletion in MCF10A cells significantly upregulated IKK-dependent p65/RELA translocation following TNFα stimulation compared to normal cells (Figure 7B). Inhibition of IKK signalling also prevented morphological changes associated with EMP (Figure 7C). JAM3 depleted RPE-1 cells also exhibited higher maximal levels of p65/RELA nuclear translocation in living cells stimulated with TNFα (Figure 7D). Thus increased IKK-NFĸB signalling is a consequence of JAM3 depletion across cell types.

During EMP we also observed extensive phosphorylation on FOXO1, IRS1, IRS2, PI3KR2, PHLPP2, RPS6KA1, RPS6KB1, PRKAA1/AMPKalpha1, PRKAB1/APMKβ1, PRKAB2/AMPKβ2, PRKAG2/AMPKgamma2, RAPTOR, and TSC2. These events are largely consistent with downregulation of INS-AKT signalling by JNK signalling, and upregulation of catabolic promoting AMPK activation. For example, we noted significantly increased phosphorylation of JNK target site Ser350 (Table S8), which suppresses signalling IRS2 function (Solinas et al., 2006). Moreover, we observed activating phosphorylation on AMPK1alpha (Thr183), which is a target of MAPK3K/TAK1 (Wang et al., 2021) (Table S8), and a simultaneous increase on the activating AMPK target site on TSC2 – which results in inhibition of mTOR signalling downstream of Insulin (Inoki et al., 2003).

Taken together we conclude that upregulation of the NFKB, JNK, and AMPK stress response pathways, and downregulation of INS-IRS1/2 signalling occurs during EMP (Figure 7D). We propose that pro-inflammatory/catabolic states may be a hallmark of EMP.

### Upregulation of the EMP network predicts mesenchymal cancers

To understand if the upregulation of EMP network is a common feature of mesenchymal subtypes across cancers, we again examined the total protein abundances of 12,197 unique proteins across 375 unique cancer cell lines from the Cancer Cell Line Encyclopaedia (CCLE) (Nusinow et al., 2020). We identified 56 lines that where VIM expression was high (>1.5) and thus considered mesenchymal, or low (<1.5 VIM) could be considered epithelial (Table S8). Of the 260 proteins of the EMP network (Figure S6), the levels of 180 were quantified reproducibly in the 56 lines (Figure 8A). The levels of these 180 proteins are sufficient to subdivide a subset of cell lines into ‘Epithelial’ (blue points in Figure 8B) versus ‘Mesenchymal’ subtypes confirmed by the high-expression of VIM (red points in Figure 8B) and/or low expression of E-Cadherin/CDH1. Meaning, that upregulation of IKK-NFĸB and INS-IRS1/2-AKT proteins signalling is a signature of epithelial and mesenchymal cancer lines in general across different genetic backgrounds.

**Figure 8.**
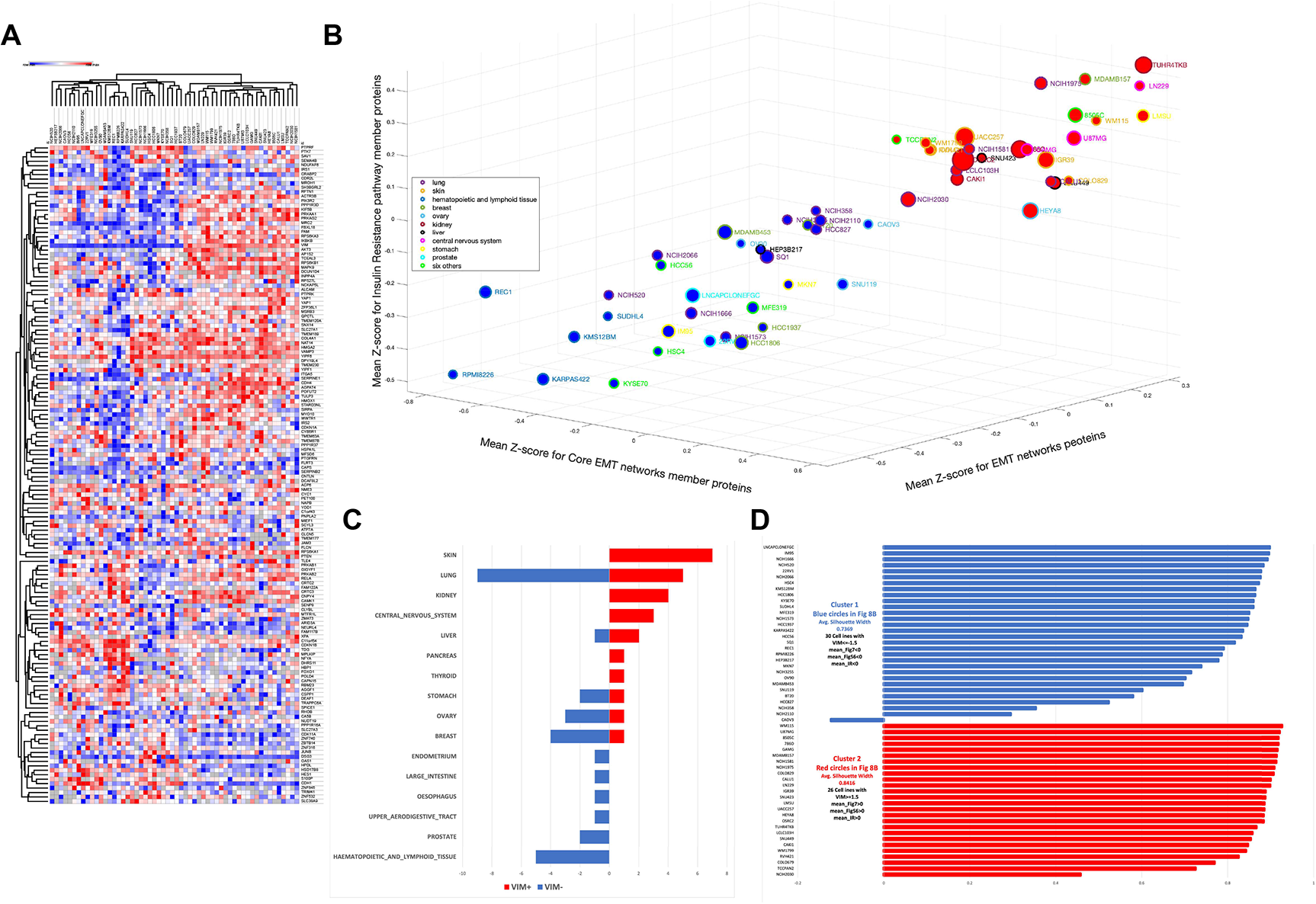
The EMP network is conserved across cancers. A) the hierarchical clustering results for 56 cell lines (columns) based on total protein intensity profiles of the Table SZ1 member proteins available in CCLE proteomics database. According to CCLE proteomics data, 26 of these 56 cell lines had z-score≥ 1.5 for VIM and average z-scores>0 when calculated among Fig 7 members (mean_Fig 7), Fig S6 members (mean_FigS6) or IR pathway members (mean_IR); the other 30 cell lines had z-score≤-1.5 for VIM and average z-scores<0 for mean_Fig 7, mean_FigS6 and mean_IR; **B)** in a space spanned by average z-scores for Fig 7 members, Fig S6 members and IR pathway members from CCLE database, the 56 cell lines in A) with positive (red) vs negative (blue) VIM values were separated in to two non-overlap groups (larger nodes indicates larger absolute value for intensity z-scores of VIM); **C)** the distribution of silhouette scores for each of the 56 cell lines as calculated from the vector of [mean_Fig7, mean_FigS6, mean_IR]; **D)** a bar chart summarizes the total protein intensity z-scores for CDH1 (yellow), JAM3 (blue) and VIM (green) from CCLE proteomics database for 60 cell lines. All cell lines had positive z-scores for VIM (green), negative z-scores for JAM3 (blue), and 51 of these 60 cell lines had negative z-scores for CDH1 (yellow), which shows that the depletion of JAM3 is non-exclusive with the depletion of CDH1. The cell lines were sorted by the VIM values.

The separation of epithelial and mesenchymal lines was further improved when considering the expression 26 ‘Insulin Resistant’ pathway proteins in the core EMP network (Figure 8B). When comparing a cluster of 26 CCLE cell lines with significantly high (z-score>1.5) VIM protein intensity with a cluster of 30 cell lines with low (z-score<-1.5) VIM intensities, the vector of mean value for members of the core EMP network generated silhouette values of 0.74 for low VIM cluster and 0.86 for high VIM cluster (Figure 8C). These silhouette values indicate good within cluster cohesion and between cluster separation. In fact, the vector of [avgCore, avgEMP, avgIR] is as effective as VIM and CDH1 expression when it comes to separating these 56 cell lines into two groups. (0.76 for low VIM cluster and 0.95 for high VIM cluster). (Figure 8C) The expression of the inflammatory-insulin EMP network was particularly upregulated in skin, kidney, and nervous system cancers, suggesting these cancers are almost essentially all mesenchymal in nature. In contrast, cancer such as breast, lung, ovary, liver, and stomach are more heterogeneous (Figure 8B).

### Inhibition of Kinesin-1 activity restores insulin signalling and inactivates FOXO3

We previously observed that KIF5B RNAi or Kinesin-1 inhibition by RBL prevents EMT, and recues changes in MTNO, by JAM3 depletion and/or TGFβ stimulation (Figures 5 and 6). To determine how Kinesin-1 can affect cell fate and shape determination, we quantified protein and phosphoprotein levels in JAM3 depleted cells (Mesenchymal) during chemical or genetic inhibition of Kinesin-1 (Epithelial-like) (Table S9). We only considered peptides increased by more that two fold with statistically significant (p<0.05) increases. We generated a network of Kinesin-1 regulated proteins using PRESTIGE (Figure 9A). Inhibition of Kinesin-1 notably downregulated pro-inflammatory signalling including the protein levels of TNF receptors TNFRSF10B, TNFRSF12A, LTBR, IKKβ (Table S12). Kinesin-1 inhibition also downregulated phosphorylation of IKKβ, SQTM1, and HMGA2 (Table S12). Moreover, multiple phosphorylation on inhibitory sites on IRS1 (Ser 527, Thr530, and Ser1101) (Copps and White, 2012) were significantly downregulated in JAM3 depleted cells treated with Kinesin-1 compared to JAM3 depleted cells alone (Table S13). Thus, we conclude that Kinesin-1 activity during EMP is largely responsible for engagement of pro-inflammatory signals which can then suppress insulin signalling (Figure 9B).

**Figure 9.**
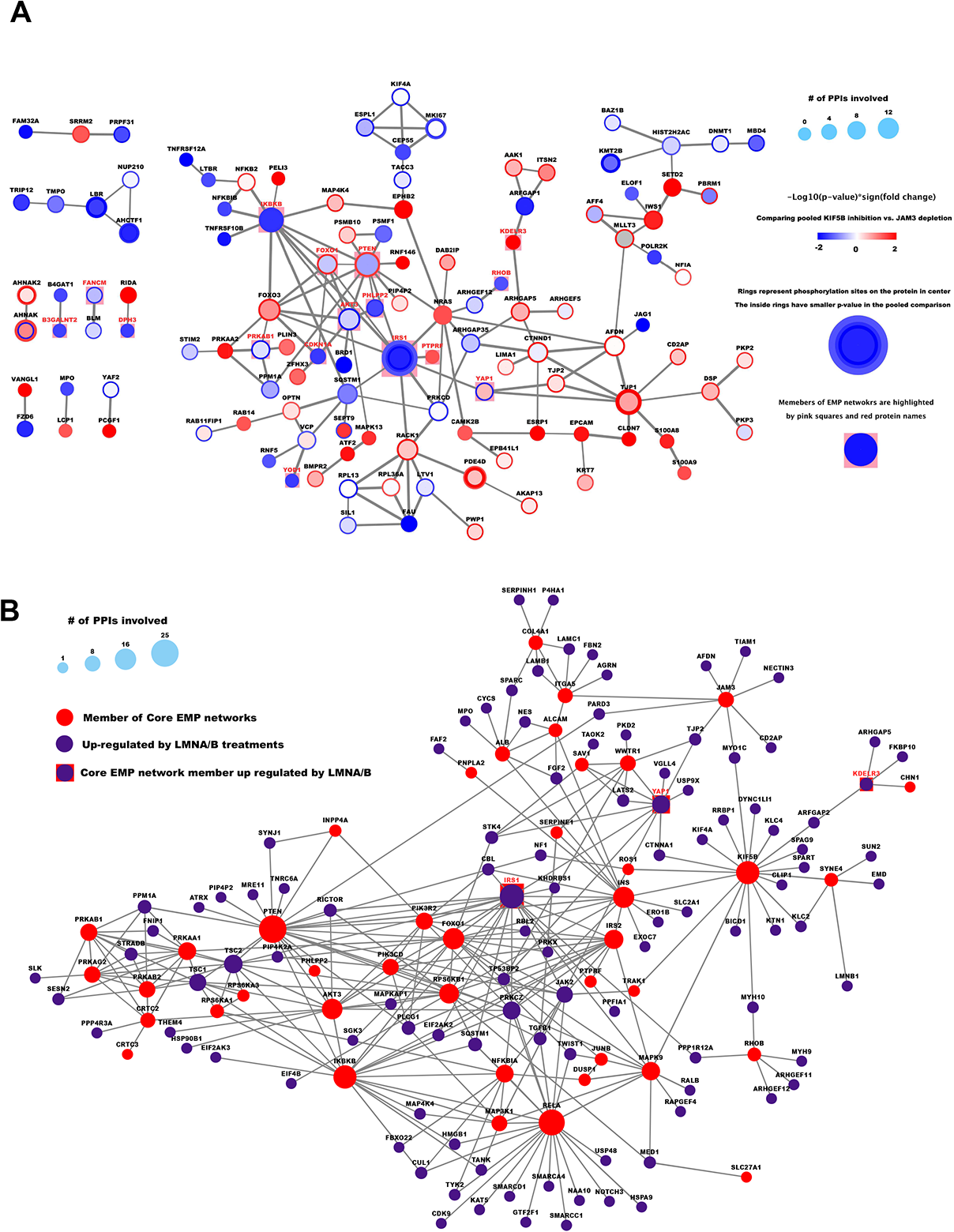
Protein subnetworks responding to KIF5B and LMNA/B treatment interacts with the EMP networks. (A) A subnetwork of 119 proteins and 153 PPIs modelled the responses to KIF5B treatment. In the pooled comparison between the proteomic abundances of KIF5B treated vs. JAM3 depleted cells, the scaled abundance of 190 phosphorylation sites and 105 total proteins had p-value≤0.05 from student’s t-test and more than 2-fold changes in either direction, including 96 phosphorylation sites and 44 total proteins that are up-regulated upon KIF5B treatments. These significantly changed sites and proteins are associated with 260 unique proteins and according to STRING database, 119 of these 260 proteins involved in 153 high confidence PPIs (confidence score ≥ 0.7). Each protein is represented by one round node and each significantly changed phosphorylation site is represented by one ring around the node for its master protein. The colour of a node of ring is proportional to the −log10 transformation of p-values from the pooled comparisons on KIF5B vs. JAM3 depletions, with red color denote up-regulation upon KIF5B treatment. The size of a node is proportional to the number of PPIs it involved in the figure. 12 of the 115 proteins in this figure are the members of the EMP network and are highlighted by a pink square in the back of the round node. (B) A subnetwork of 134 proteins and 294 PPIs modelled the interaction between the core EMP networks and subnetworks upregulated by change in nuclear shape. In the pooled comparison between the proteomic abundances of (LMNA-LMNB-SYNE4) vs. control, a group of 94 proteins are highlighted as purple circles. These 94 proteins are associated with 52 total proteins and 67 phosphorylation sites whose scaled abundance vectors presented p-value≤0.05 from student’s t-test in the pooled comparisons and were up-regulated in (LMNA-LMNB-SYNE4) condition. 3 of the 94 proteins are members of core EMT network (purple circles with red squares in the back), and overall, these 94 proteins also involved in 191 high confidence PPIs with 40/48 members of core EMT networks (red circles if not within the group of 94). High confidence PPIs within two proteins of the same color were also included to make a network of 136 unique proteins (91 purple circles, 3 purple circles with red squares, 37 red circles interacting with purple circles, 5 red circles only interacting within this group of 5) and 295 PPIs. Each protein is represented by one round node. The size of a node is proportional to the number of PPIs it involved in the figure.

In contrast, after Kinesin-1 inhibition we see significant upregulation of CLDN7, EPCAM, ESRP1, and KRT78 protein. Additionally, we observed upregulation of phosphorylation on regulators of MTs (q=3.9805E-07), Adherens junction (q=4.7900E-05) and Tight Junction (q=0.0004947) including CTNND1, DSP, KRT7, KRT10, LIMA1 MLLT3, PKP2, PKP3, TJP1, and TJP2. (Table S13, S14) We interpret these molecular remodelling events as evidence of ‘epithelization’ of cells that occurs when Kinesin-1 is inhibited (Figure 9A, TableS12).

### AKT signalling is restored by inhibiting Kinesin-1 during EMP

Notably, following Kinesin-1 inhibition in JAM3 depleted and/or TGFβ treated cells we observed significant increases in the phosphorylation of FOXO3 at Ser253. Ser253 is an AKT target, that when phosphorylated promotes binding of 14-3-3 binding proteins which restrain and inactivate FOXO3 in the cytoplasm (Table S11) (Tzivion et al., 2011). These data suggested to us that FOXO3 phosphorylation and nuclear sequestration downstream of activated Insulin-AKT may occur during suppression of EMP. To confirm this idea, we measured levels of nuclear FOXO3 in wild-type, JAM3, TGFβ stimulated cells. In only wild-type cells was FOXO3 predominantly cytoplasmic (Figure S7A-C) which infers the AKT signalling (AKT1 or AKT3) is activated in these cells. In both JAM3 and TGFβ treated cells, FOXO3 nuclear levels increased (Figure S7A-C). This observation is markedly consistent with our idea that EMP coincides with downregulation of INS-AKT signalling. Treatment of either JAM3 and/or TGFβtreated cells with the Kinesin-1 inhibitor suppressed FOXO3 translocation (Figure S7A). Although nuclear FOXO3 levels correlated across population with average nuclear morphology, this was not the case at the single cell levels (Figure S7B). We conclude that inhibition of AKT signalling and activation of FOXO3 is a hallmark of EMP.

### Nuclear shape alterations affect effectors of inflammatory and insulin signalling but not core enzymes of the EMP network

Taken together our data demonstrates that Kinesin-1 activity is required for both upregulation of EMP, and coincident changes in LMNA levels and nuclear shape. However, to this point we have been unable to determine if changes in nuclear shape are also causal to EMP. To determine if changes in nuclear lamina morphology drive EMT, we performed mass spectrometry analysis of LMNA, LMNB, and SYNE-4 depleted cells (Table S11). These are cells with gross alterations in nuclear shape but where JAM3 or Kinesin-1, and thus microtubule organization, have not been deliberately targeted. We considered these as a combined group which we use to compare to cells with ‘normal’ nuclear morphology (Methods). This increases our statistical power and facilitates the identification of proteins whose levels are affected by *any* change in nuclear shape. Changes in nuclear shape coincided with upregulation of 337 total proteins and 405 phosphorylation sites corresponds to changes in 623 unique proteins (p-value<0.05). There are 804 high confidence (confidence score≥0.7) PPIs involving 385 of 623 unique proteins (an average of 2.0883 high confidence PPIs per protein; Table S14) These interactions define a network of proteins affected by alterations in nuclear shape (Figure 9C).

Among the proteins/phosphopeptides that were upregulated by changes in nuclear shape only IRS1, YAP1, and KDELR3 were also altered following induction of EMP by JAM3 depletion or TGFβ stimulation (Figure 9B, Table S14). Thus, clearly changes in nuclear shape alone do not explain the signalling state caused by JAM3 depletion/TGFβ stimulation.

While nuclear shape proteins form a network amongst themselves, we asked if nuclear shape regulated proteins might interact with the core EMP network of 48 proteins that is upregulated by JAM3 depletion and/or TGFβ stimulation (Figure 7) and conserved across mesenchymal cancers (Figure 8). 94 of 623 nuclear shape proteins interacted with 40/48 members of the core EMP network for a total of 191 high confidence PPIs involving one nuclear shape regulated protein and one member of core EMP network (Figure 9B). Interestingly, this sub-group of 94 proteins was a poorly connected subset within the 623 nuclear shape responding proteins themselves, as only 69 of these 94 proteins involved in 88 high confidence PPIs within the group. However, the integration with the core EMP network increased the general connectivity of 94 member sub-group (the number of PPI per protein increased from 0.9362 within the group of 94 to 2.0319 when combined with the core EMP networks). The average number of high confidence PPIs for the group of 40 members within the core EMP network also increased substantially from 1.9 within the core EMP network (36 of the 40 members involved in 76 high confidence PPIs) to 4.775. Typically, this was the result of core EMP proteins gaining multiple individual interactors and becoming ‘hubs’ for different complexes (Figure 9C).

These observations suggests that nuclear shape changes engage similar pathways as JAM3 depletion and/or TGFβ stimulation, even though the specific proteins regulated are not identical. For example, when integrating nuclear shape regulated proteins, the transcription factor RELA acted as a “hub” and gained 16 interactors, none of which have high confidence interactions within themselves. Of these 3/16 (TWIST1, NAA10, and HSPA9) do not have any high confidence PPIs within the group of 94. Notable amongst these are upregulation of canonical EMT inducers such as TGFβ, FGF2 (Lee and Kay, 2006). Changes in nuclear shape alone also resulted in increased phosphorylation of the pro-EMT transcription factor Twist-1 in Ser68, which stabilizes Twist-1 (Hong et al., 2011). Similarly, proteins involved in canonical INS-AKT, IKK-NFĸB, and Hippo-YAP/TAZ, all interacted with proteins affected by nuclear shape. Thus, these data suggest that changes in MT and Kinesin-1 activity upregulate hub proteins such as RELA, IKBKB, IRS2, and TAZ, but EMP is also driven by changes in nuclear shape upregulate key effectors of these hubs (Figure 9C).

We reasoned that if the 94 proteins that are upregulated by nuclear shape shapes are modifiers of canonical signalling pathways that drive EMP their levels should be affected in JAM3KD/TGFβ – albeit potentially in a modest fashion. For the 94 nuclear shape regulated proteins that interacted strongly with the core EMT network, we detected changes in protein levels changed for 52/94 proteins and measured phosphorylation for 67/94 of them following nuclear shape changes. Of these, 5/52 total proteins and 20/67 phosphorylation sites were significantly upregulated upon JAM3KD/TGFβ-stimulation as predicted. Thus, the most profound similarity between JAM3KD/TGFβ-stimulation and nuclear shape changes alone was in the regulation of protein phosphorylation versus protein levels (Table S14). For example, we detected upregulation of phosphorylation on the pro-EMT transcription factor TWIST Ser68 TSC2 Ser1411, SQSTM1 Serine 24 in JAM3 depleted and TGFβ stimulated cells, as well as in cells with altered nuclear shape. Thus, we conclude that changes in nuclear morphology mechanistically contribute to the engagement of EMT through modifying the activity of canonical signalling pathways that are upregulated in a Kinesin-1 dependent fashion following JAM3KD/TGFβ-stimulation. Some of these events, including regulation of Twist-1 are likely key to long term stabilization of EMP.

### JAM3 expression is associated with good prognosis

To validate JAM3’s role as a tumour suppressor in patients, we performed survival analysis using JAM3 expression and disease-specific survival in different subgroups of breast cancer. We observed that in all subgroups with elevated levels of JAM3 (Luminal A, ILC, and Normal-like intrinsic subtypes), JAM3 expression is correlated with favourable prognosis (Figure 10A). Increased JAM3 expression also predicts relapse-free survival after treatment in patients with Luminal A type tumours (Figure 10B). Thus JAM3 is a *bona fide* tumour suppressor in breast cancer patients.

Given the ability of JAM3 depleted MCF10A cells to invade 3D matrices, we next sought to determine if JAM3 altered the metastatic potential of breast tumour cells in patients. Although all forms of metastasis may not involve entry in a mesenchymal state (Fischer et al., 2015; Somarelli et al., 2016; Zheng et al., 2015), the metastasis of cells to stiff tissues such as bone likely requires matrix remodelling proteins such as MMPs whose levels are elevated following JAM3 depletion (Figure 3G)(Esposito et al., 2018; Kang et al., 2003). Indeed, we observed a 1.6-fold increase of bone metastasis in cases with JAM3 loss compared with copy number neutral cases (Figure 9C), which suggests that cells with decreased levels of JAM3 have great invasive potential *in vivo*.

We reasoned that if KIF5B-SYNE4 mediated linkage of MTs to the nucleus is essential for nuclear shape changes following loss of JAM3, KIF5B and/or SYNE4 may be oncogenes. Indeed tumours with low JAM3 expression have high levels of KIF5B and SYNE4, whereas cells with high levels of JAM3 have low KIF5B and SYNE4 expression (Figure 10D). Moreover, increased KIF5B and SYNE4 copy numbers were observed in tumours with low JAM3 expression (Figure 10E). Finally, increased KIF5B and SYNE4 expression predicted shorter disease-specific survival periods (Figure 10F). Thus KIF5B and SYNE4 are oncogenes that antagonize the actions of the JAM3 tumour suppressor in patients.

## Discussion

Our results show how stereotypical changes in cell/nuclear shape and cytoskeletal organization promotes epithelial-mesenchymal plasticity (EMP) by establishing a conserved signalling state. We thus link changes in cell shape to alterations in signalling network dynamics and cell fate decisions that drive diseases such as cancer. Importantly, our data provide a mechanistic basis for how transient dysregulation of epithelial tissue morphogenesis, such as caused by defects in cell-cell adhesion formation (Kobielak and Fuchs, 2006; Perez-Moreno and Fuchs, 2006; Vasioukhin et al., 2001) and/or loss of apicobasal polarity (Bilder et al., 2000; Bilder and Perrimon, 2000; Martin-Belmonte and Perez-Moreno, 2011; Pearson et al., 2011; Zhan et al., 2008), or even changes in environment (Greenburg and Hay, 1982), can trigger EMP, stabilize transcriptional programmes, and accelerate oncogenesis (Cordenonsi et al., 2011; Dongre and Weinberg, 2019). More generally they suggest that any mechanical and soluble cue that drives EMP may do so in part by driving changes in MT organization and nuclear shape. In fact, we predict similar morphological and signalling events may happen in other cell types during fibrotic, metabolic, and neurodegenerative disorders.

Using multiple ways to genetically and chemically induce EMP, we show that the EMP is characterized by radial MT organization and abnormal nuclear morphology. We propose these morphological changes are coupled to changes to upregulation of an EMP network comprised of INS-IRS-FOXO, IKK-NFKB, and YAP/TAZ pathways leading to the morphogenesis of pro-inflammatory, insulin-resistant, pro-invasive mesenchymal cells.

Based on these data and our previous observations (Sero et al., 2015), we suggest that EMP occurs during a number of events that may happen in a coordinated manner. We propose cell-cell adhesions, and in particular JAM3, are essential to capture and stabilize MTs, or even nucleate the polymerization of new MTs at the cell cortex. In these epithelial cells, MT-directed Kinesin-1 activity acts largely to position the nuclei through interactions with SUN-KASH complexes, and in particular SYNE-4. (Roux et al., 2009; Wilson and Holzbaur, 2015; Wu et al., 2018). Trigged by a mechanical cue – such as change in cell-cell adhesions architecture (i.e. JAM3 depletion) – MTs detach from the cell-cell adhesion complexes, and a “collapse” of MTs into a radial structure. This model is consistent with data demonstrating that JAM-A, which is highly related to JAM3 has been shown to bind the polarity proteins PARD3 (Ebnet et al., 2001; Itoh et al., 2001), and PARD3 is critical for recruitment of MT nucleators such as gamma-tubulin to the apical membrane (Feldman and Priess, 2012). MT collapse into a radial array leads to concomitant alteration in the dynamics of Kinesin-1 cargo binding and trafficking. In particular we propose Kinesin-1 transports cargo to the leading edge to promote EMP. Indeed, Kinesin-1 activity has been previously shown to be essential for transporting cargoes to the leading edge of invasive mesenchymal cells which are essential for migration and morphogenesis (Marchesin et al., 2015; Wang et al., 2017). Kinesin-1 is also required for TGFβ driven EMP (Moamer et al., 2019).

During EMP we propose that Kinesin-1 activity and cargos are especially important to upregulate stress response/pro-inflammatory signalling, especially that of JNK and IKK-NFĸB pathways (Figure 9A). A likely link between Kinesin-1 activity and JNK activity is likely made by JIP (JNK interacting proteins) proteins, which simultaneously bind Kinesin-1 and JNK activators (Kelkar et al., 2005; Marchesin et al., 2015; Schulman et al., 2014; Sun et al., 2011; Verhey et al., 2001). Indeed we observed that significantly increased phosphorylation of MAPK8IP3/JIP-3 during EMP (Table S8).

Though the role of JIP-3 phosphorylation is poorly understood, our data strongly suggests that Kinesin-1/JIP-3 mediate the assembly of large complexes which promote JNK activation. JNK in particular has been extensively implicated in regulating mesenchymal morphogenesis and behaviours such as migration and invasion (Kalli et al., 2021). IKK-NFĸB pathways have also been implicated in EMP by ourselves and others (Alameda et al., 2011; Alameda et al., 2016; Goktuna et al., 2018; Sailem and Bakal, 2017; Sero et al., 2015). Increases in JNK activity during EMP in particular are likely for downregulation of Insulin signalling, and upregulation of AMPK signalling a number of different means, in particular via inhibitory IRS1/2 phosphorylation by JNK kinases (Copps and White, 2012; Garg et al., 2021; Holzer et al., 2011; Solinas et al., 2006; Solinas et al., 2007) (Figure 9B).

One significant role of Kinesin-1 during EMP appears to be deforming the nuclei, which engages a sub-network of effectors of core EMP components (Figure 9C). A role of Kinesin-1 in nuclear morphogenesis has been found in numerous organisms (Bone et al., 2016; Fridolfsson et al., 2010; Fridolfsson and Starr, 2010; Lottersberger et al., 2015; Roux et al., 2009; Simon and Wilson, 2011; Splinter et al., 2010; Tanenbaum et al., 2011; Wilson and Holzbaur, 2015; Wu and Kengaku, 2018; Wu et al., 2018). Moreover, in *Drosophila* Kinesin-1 acts together with JIP proteins to position nuclei (Schulman et al., 2014). Determining whether these nuclear deformations that drive EMP in manner distinct from MT reorganization and Kinesin-1 activity is technically challenging. However, we show that depletion of LMNA, LMNB, or SYNE-4 – which results in nuclear deformation independent of Kinesin-1 depletion − upregulates a sub-network of proteins that show average interactions amongst themselves but form a large number of interactions with the core EMP network (Figure 9C). In many cases the components engaged by nuclear deformation appear to be modifiers of canonical signalling components such as PTEN, RELA, and TAZ (Figure 9B). In particular, changes in nuclear shape upregulate the levels and/or phosphorylation of established pro-inflammatory, pro-EMP proteins such as TNFαreceptors and TWIST-1 (Figure 9C).

Our data strongly points to a role for both YAP and TAZ transcription factors in promoting EMT. Depletion of either YAP or TAZ partially suppresses mesenchymal phenotype in 2D and 3D models. Based on the signalling pathways that are engaged during EMP, how YAP/TAZ become activated during EMP is not obvious. But we show YAP/TAZ activation is clearly dependent on MT reorganization and/or Kinesin-1 activity (Figure 5). One simple explanation is that increased formation of focal adhesions (FA) in mesenchymal cells results in increased YAP/TAZ activation (Sero and Bakal, 2017). Such a model would be consistent with our observation that YAP in particular is responsible for modelling of the ECM during EMP in 3D models. Perhaps surprisingly, YAP/TAZ activation does not appear to correlate with changes in nuclear shape (Figure 4H). In fact, changes in nuclear shape appear to partially dependent on YAP/TAZ during EMP (Figure S5F, S5G). Thus, we propose that YAP/TAZ activation – perhaps driven by FAs during mesenchymal morphogenesis – further ‘stabilizes’ mesenchymal phenotypes by altering nuclear architecture. These changes in appear to activate pro-EMT TFs such as Twist-1.

Taken together our finding suggest that EMP might be initiated in the short-term by the simultaneous engagement of multiple canonical ‘generic’ pathways such as AKT, NFĸB, and Hippo signalling. But that this signalling state is stabilized by changes in nuclear shape. In the case of breast epithelial cells, large scale changes in cell-cell adhesion and MT reorganization can activate an EMP network – i.e. during transient fluctuations in soluble or mechanical cues. But that prolonged activation of this network leads to nuclear deformation – which sustains a signalling state that sustains EMP. The idea that changes in nuclear shape sustains an EMP state initially trigged by changed in cell-cell adhesion and MT organization is very consistent with our previous finding that while TNFα can promote NFĸB activation in both epithelial and pre-mesenchymal cells in the short term – only cells with specific cell and nuclear shapes undergo differentiation to a more mesenchymal state (Sero et al., 2015).

A pro-inflammatory insulin-resistant state is a hallmark of many different types of diseases including cancer (Grivennikov et al., 2010). In this study we show that the network of upregulated proteins is common across multiple cancer sub-types and largely distinguishes epithelial and mesenchymal forms of the disease (Figure 8). Moreover, such states are common to diabetes, neurodegenerative disorders, and infection driven syndromes – including severe COVID-19 (Lambert and Weinberg, 2021; Nieto et al., 2016; Pandolfi et al., 2021). This raises the possibility that cell shape changes which results in MTNO, and/or alter Kinesin-1 activity drive a common state which leads to different disease based on the cell of origin.

Our work thus clearly opens up stratification and therapeutic opportunities for both cancers and potential other diseases underpinned by EMP. For example, we show that in breast cancers JAM3 loss is a common event, but is also frequently lost in other mesenchymal cancers; consistent with previous reports (Zhou et al., 2019). We would thus predict that inhibition of Kinesin-1 activity would be one mechanism by which to limit EMP, and ultimately disease aggressiveness as “differentiation” therapy. Indeed our work provides dozens of new targets for such work (Pattabiraman et al., 2016; Pattabiraman and Weinberg, 2016).

## Author Contributions

M.A.-G, Z.Y., Y.Y., and C.B. conceived the research. M.A.-G designed and performed all the experiments, developed image analysis, and analysed all the experimental data. Y.Y. performed the bioinformatics analysis on clinical samples. T.I.R, and J.S. performed the proteomics sample analysis and generated the proteomics data. Z.Y., L.W. and S.T.C.W developed PRESTIGE and performed all statistical cell biology analysis. R.R. executed the sub fractionation experiments. M.G.-A, Y.Z., Y.Y., and C.B. wrote the manuscript.

## Acknowledgements

We thank A. Sadok (Cancer Therapeutics, ICR, London, UK) for technical help in preliminary bottom to top invasion assays. We also thank J. Sero (Biology and Biochemistry Department, Bath University, UK) for initial help to quantify donut nuclei on Acapella. This project was initiated by Y.Y. in the laboratory of Florian Markowetz (Cancer Research UK Cambridge Institute). C.B. was supported by a Stand Up to Cancer UK Programme Foundation Award (C37275/1, A20146) and a CRUK/EPSRC Multidisciplinary Research Award (NS/A000062/1). Y.Y. acknowledges funding from Breast Cancer Now (2015NovPR638) and CDMRP Breast Cancer Research Program Award BC13205 to support this work. Y.Y. acknowledges additional support from Cancer Research UK Career Establishment Award (CRUK C45982/A21808), CRUK Early Detection Program Award (C9203/A28770), CRUK Sarcoma Accelerator (C56167/A29363), CRUK Brain Tumour Award (C25858/A28592), Rosetrees Trust (A2714), Children’s Cancer and Leukaemia Group (CCLGA201906), NIH U54 CA217376, NIH R01 CA185138, European Commission ITN (H2020-MSCA-ITN-2019), and The Royal Marsden/ICR National Institute of Health Research Biomedical Research Centre. Z.Y., L.W. and S.T.C.W are supported by the National Institutes of Health (NIH R01CA238727, NIH U01CA253553, NIH R01CA244413), the T.T. & W.F. Chao Foundation, and the John S Dunn Research Foundation.

## EXPERIMENTAL MODEL AND SUBJECT DETAILS

### Cell lines and Cell Culture

MCF10-A cells were obtained from ATCC. They were cultured in DMEM/F12 supplemented with 5% Horse Serum, 10 μg/ml insulin, 20 ng/ml epidermal growth factor, 100 ng/ml cholera toxin, 500 ng/ml hydrocortisone and 100 mg/ml penicillin/ streptomycin. Cells were used between passages 4 and 15. hTERT-RPE1 cells were obtained from ATCC and cultured in DMEM, 10% heat-inactivated FBS and 100 mg/ml penicillin/ streptomycin.

All cell lines were confirmed to be mycoplasma-negative (e-Myco Mycoplasma PCR Detection Kit, iNtRON Biotechnology). Passage was carried out using 0.25% trypsin-EDTA (GIBCO) followed by centrifugation (1000 rpm, 4 min) and resuspension in complete medium. Cell counting was performed using Countess automated cell counter with trypan blue exclusion (Thermo).

### Clinical samples

Primary frozen breast tumors were collected and stained independently in different laboratories contributing to the METABRIC consortium. All patient specimens were obtained with appropriate consent from the relevant institutional review board. All METABRIC samples used in this study (n=1026) included H&E images and molecular data passing quality control. Control DNA from match normal breast tissues was available for 473 samples and RNA from 144 of these individuals. All histopathological images were scanned with a ScanScope TX scanner (Aperio Technologies Inc.) providing images at 200-fold magnification. DNA and RNA were extracted from each primary tumor specimen and subjected to copy number and genotype analysis on the Affymetrix SNP 6.0 platform and transcriptional profiling on the IlluminaHT-12 v3 platform (Illumina_Human_WG-v3).

### Pathological image analysis

To facilitate objective extraction of nuclear morphological features for tens of thousands of cells in a histological section, we performed automated image analysis on 1,026 H&E images of breast primary tumour whole sections using a previously described pipeline (Yuan et al., 2012). In brief, this pipeline segmented and classified nuclei into cancer, lymphocyte, and stromal cells based on their morphological features. To remove batch effects as a result of H&E slides batch processing, we normalized scores after subtracting median value of the batches by which images H&E were stained and scanned.

Following cell nuclei segmentation and classification, six nuclear shape features were used for each cancer cell nucleus, which include acircularity (*acirc*) and Hu’s first moment or I1 (*I1*) both measuring shape irregularity. While acircularity is mathematically defined as the fraction of object area outside of a circle of the same size of area as the object, *I1* is equivalent to the moment of inertia and considers not only the shape but also the distribution of mass in the object. Hu’s first moment or I1, known as the moment of inertia in mechanics, has been used to define an object’s resistance to rotation in physics. In 2D images, pixel density is equivalent to physical mass within an object.

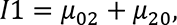

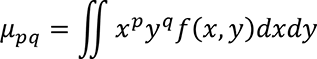

where *f(x,y)* is the image segment. This morphological feature, when applied to nucleus image, represents nuclear irregularity not only in shape but also in the nuclear mass distribution.

### Construction of a hybrid network

Significant associations followed by FDR correction between 18 morphological scores and RNA expression of 1000 cis-driven genes were used to construct a large network. Small network module shown in Figure 1B was constructed by removing associations with q-value greater than 0.005. Degree of connectivity was assessed for each morphological score, based on their connections with genes within the large network as well as the small network module.

### RNAi transfections and drug treatments

RNAi experiments were performed by using the siRNA oligonucleotides from Dharmacon (see Supplemental Table S5 for the sequences). Cells were reverse-transfected in either 384-well, 96-well, or 6 wells plates, T25 or T75 flasks either with single siRNAs or combinations thereof. 2 days later cells were either fixed or harvested for the bottom to top invasion assays, organotypic experiments, mRNA extraction, protein extraction or mass spectrometry.

The majority of experiments were performed in 384-well Cell Carrier plates using MCF10A cells by using an in lab optimized screen protocol. Two siRNAs targeting *ECT2* and *PLK1* at several concentrations were used as transfection controls in order to test the efficiency of the transfection (data not shown). Cell density and transfection times were assessed. The output of multinucleated cells for cells lacking *ECT2*, cell death for cells lacking *PLK1* and cell number was used to choose the best conditions. In order to transfect cells in the other 3 formats, all the reagents and cells were scaled up from the 384-well format according to the surface area and concentration in order to keep the same conditions across all the culture conditions.

All the siRNAs were used at a final concentration of 20nM. Briefly, 5uL per well of OptiMEM (GIBCO) containing 40 nL/well Lipofectamine RNAiMAX (Invitrogen) were mixed with 5uL per well of OptiMEM containing also 40 nL/well of one single siRNA or combinations. The mix was incubated for 30-40 min at room temperature to allow the siRNA-Lipofectamine RNAiMAX complexes to form. Then, 30 uL of 4/3X Transfection media (to give 1X final concentration of serum and growth factors) containing cells were plated on top of the complexes at 3×10^4^ cells/ml. For the experiments regarding YAP and TAZ immunofluorescence a density gradient of mock-transfected cells (375, 750 and 1500 cells per well) was plated to use as a density regression curve (S9L and S8M). siRNAs targeting *YAP1*, *TAZ, LMNA, and LMNB1* were also used to confirm gene knockdown and antibody specificity. All the 2D experiments were performed by using at least 4 wells per condition per experiment, and at least two independent biological replicates were performed for all the imaging experiments.

Rose Bengal Lactone was used at 20uM or 40uM for 48 hours. The broad-spectrum MMP inhibitor GM6001 was used at 10uM for 24 hours before fixing the bottom to top invasion assays in collagen. Transforming Growth Factor-β1 was used at 5ng/mL for 48 hours (IF) or 2h (Proteomics). Stimuli with Recombinant Human TNF-α Protein was performed at 10ng/mL for 30min (IF) or 2h (Proteomics). BAY 11-7082 was used at 10uM for 2h.

### Fixation and immunofluorescence in 2D

Following two days of incubation, cells were fixed either in pre-warmed 4% formaldehyde (ThermoScientific) in PBS for 15 min at room temperature (RT) or with cold MeOH for 10 min (anti-JAM3). After fixation, cells were washed three times in PBS and then permeabilised in 0.2% Triton X-100/PBS solution for 15 min at RT (this step was not done for MeOH fixed samples). Following three washes in PBS, cells were blocked for 1 h in 2% bovine serum albumin (BSA) (Sigma)/PBS solution at RT. When using both mouse and rat primary antibodies in the same sample, sequential immunostaining was performed to avoid any antibody cross-reactions. Typically co-immunostaining with a mouse, rat and rabbit antibody was used. After the Block step, BSA was removed and the desired mouse primary antibody was added in Antibody solution (0.5%BSA/0.01% Triton X-100/PBS) at the indicated dilutions: JAM-C (Santa Cruz, 1:200), YAP (G6) (Santa Cruz, 1:200), YAP/TAZ [67.3] (Santa Cruz, 1:1000). All the primary antibodies immunostaining was performed overnight at 4 °C. Then cells were washed three times in PBS and incubated with a goat anti-mouse antibody 1:1000 in Antibody solution for 2h at RT. Cells were washed three times in PBS and incubated with a rat anti-tubulin α antibody (Bio Rad, 1:1000) and an anti-rabbit primary antibody when applied, for 2 hours at RT. The anti-rabbit primary antibodies were used at the indicated dilutions: Phospho-Lamin A/C (Ser22) (Cell Signalling, 1:800), TAZ (V386) (Cell Signalling, 1:200), Anti-Lamin B1 (Abcam, 1:1000), FOXO3a (Cell Signalling, 1:100). Then cells were washed three times in PBS and incubated for 2h at room temperature with a goat anti-rat antibody and/or a goat anti-rabbit antibody or Alexa-647 phalloidin (Invitrogen) if needed. Finally, to stain nuclei, a 1:1000 dilution of DAPI (4’,6-diamidino-2-phenylindole, dihydrochloride, Molecular Probes; D1306, Molecular Probes)/PBS solution or 5 mg/ml Hoescht (Invitrogen)/PBS solution was carried out for 15 min at RT. 384-well plates were sealed for imaging with an Opera Cell:Explorer-automated spinning disk confocal microscope (PerkinElmer) in the magnification indicated in the figure legends. At least twenty fields at random positions per well of a 384-well plate were imaged.

### Bottom-Top Invasion screens in a Collagen matrix

MCF10A cells were transfected with 20nM of RNAi for 48h as described above. Then cells were trypsinized and suspended in a solution of NaOH, DMEM 5x and bovine collagen I (Purecol) at 2.3 mg/ml to a final concentration of 5 × 104 cells/500 μl. Bovine collagen neutralization at pH=7 was confirmed in all the experiments performed. One hundred microliter aliquots per well of neutralized collagen solution were dispensed into chilled Cell carrier 96-well plates followed by a centrifugation at 1000 rpm at 4°C for 5 minutes to ensure all cells were at the bottom of the plate. Then, plates were incubated at 37°C and 80% humidity for 3h to allow the collagen to polymerize. Importantly, the outer wells were also filled with sterile water or PBS to prevent gel dehydration and edge effects due to evaporation. Then 50uL of growth media 3X containing 30uM of the pan MMP inhibitor GM6001 or DMSO was added on the top to stimulate cell invasion. After 24 h of incubation at 37°C, cells were fixed at room temperature with 50 uL per well of 16% Methanol-free paraformaldehyde plus 5 μg/ml Hoechst 33258 for 24 h. Samples were run in quadruplicate for each condition and repeated twice in experiments separated by several months. Cells were stained with 1:500 phalloidin/ PBS solution overnight with a previous permeabilization step with NP40 at 0.5% at 4°C for 30 minutes. Plates were sealed and imaged with an Opera Cell:Explorer-automated spinning disk confocal microscope (PerkinElmer) and using a 20× air objective lens (NA = 0.45). Three Confocal Z slices were collected from each well at 0 μm (bottom of well), 30 μm (middle of the well) and 60 μm (top of the well) and at least 20 fields were imaged in every Z stack position in each well avoiding the edge fields.

### Invasion of Organotypic cultures in a Matrigel:Collagen matrix

The morphogenesis assay was adapted from (Debnath et al., 2003). Briefly, previously chilled eight-well culture slides (Falcon) or cell carrier 96 wells plates were coated with a thick layer (150-200uL/cm2) of a mix of growth factor-reduced (GFR) Matrigel (BD Biosciences) and neutralized Bovine Collagen I (Purecol) yielding a final collagen concentration of 1 mg/mL and a final Matrigel gel mix concentration of 5 mg/mL. The stock solutions of all the Matrigel vials used to perform the experiments of this manuscript were 10 mg/mL at least (See Supplemental Table S3 for the lot numbers used). Importantly, all the surrounding wells were also filled with water or PBS and the ECM was gelled for 3 hours in a standard 37 °C humidified cell culture incubator with a 5% CO_2_ environment. Then, a single MCF10A cell suspension (4000 cells/well for 96 wells plate or 8000 cells/well for 8 wells chambers at 2*104 cells/mL) was plated on top of the pre-polymerized Matrigel:collagen matrix in acini assay medium. This assay media is composed of phenol red free DMEM supplemented with cholera toxin, hydrocortisone, insulin and penicillin/ streptomycin at the same concentrations than the growth media and 2%horse serum, 5 ng/ml of EGF and 2mg/mL of Matrigel. Unless otherwise stated, the acini morphology data refers to the structures observed 2 days after the 3D experiment started and 4 days after the cells were transfected with siRNA. When acini were cultured for 4 days or 8 days, assay medium was replaced every 2-3 days. Thus 4 or 8 days of acini growth in the main text corresponds to 6 or 10 days after the siRNA transfection. Acini were fixed in 4% Methanol-free paraformaldehyde and 5 μg/ml Hoechst 33258 for 15 min at RT. Inmunostaining was performed with Laminin-5 Antibody (P3H9-2) (Santa Cruz, 1:1000) and anti-Fibronectin (Dako, 1:1000) in Block solution (4% BSA) plus 0,05% of Tween20 overnight at 4 degrees with a previous permeabilization step with 0.5% Triton X-100/PBS solution for 30 min at RT and blocked in 4% BSA for 2 hours at RT. Acini were washed 3 × 20 minutes in PBS and then incubated with a goat anti-mouse 568 antibody and with a goat anti-rabbit 488 antibody in block solution plus 0,05% tween20 for 1 h at room temperature. Acini were washed 3 × 20 minutes in and finally a 1:500 phalloidin 647/ PBS solution was added either to the 8-well chamber or to the 96-well plate, sealed and left overnight at 4 degrees. 8 -well chamber slides were rinsed in PBS, mounted in Vectashield (H-1000, Vector Laboratories) and the coverslip was sealed with clear nail varnish. Slides were imaged using either a Zeiss 710 LSM Confocal microscope or a Leica SP2 confocal scanning microscope (Leica Microsystems, Milton Keynes, Bucks, UK) at 20x or ×40 oil immersion lens as indicated in the figure legends. At least 400 acini were counted by eye and classified according to their shape in normal acini (round) or abnormal shape (general loss of spherical geometry and/or protrusions). See the figure legends for more details. 96 well plates were sealed and imaged with an Opera Cell:Explorer-automated spinning disk confocal microscope (PerkinElmer) and using a 20× air objective lens (NA = 0.45). Several Confocal Z slices were collected from the bottom to the top of the acini and they were summed to create a projection of the image (maximum projection). At least 9 fields were imaged per well. Importantly only the middle and symmetrical fields were imaged avoiding possible lateral interactions of the acini with the plastic sides of the culture dish.

### Cell Segmentation, shape feature extraction and analysis of 2D and 3D experiments

Image acquisition and cell segmentation was performed using Columbus high-content image analysis software. ‘Whole Nuclei’ were segmented using the Hoechst or DAPI channel.

<u>For all the 2D experiments</u>, cell bodies were segmented using the tubulin channel (Figure S9B). The cell-cell contact area (Neighbour fraction) was determined using an inbuilt Columbus algorithm ‘Cell Contact Area with Neighbors [%]’ expressed as the Percent of the object border that touches a neighbouring object. The border objects were removed from the analysed cells considering only cells completely imaged. Mitotic cells were filtered using a combination of Hoechst intensity mean and Hoechst intensity maximum and excluded of all the analysis of this study. Since several siRNAs produced holes in the nuclei of the cells, the nucleus region used for all the experiments of this manuscript was limited to a fixed region to avoid counting holes as part of the nuclei and named as ‘nuclei’ or ‘Inner nuclei’. The perinuclear region was used to measure cytoplasmic antibody intensities. To classify the cells according to their nuclei shape we used a designed Columbus script where the mean intensity and the size of the whole nuclei was calculated. The ‘donut population’ was detected by a threshold mask for Hoechst minimum intensity that detected the holes of the donut nuclei and split the whole population in donut population or non donut population (S9G). To filter the cells with low Nuclear Roundness, a threshold of Nuclear Roundness < 0.85 was established within the non donut population, hence excluding the cells having a donut nuclei. The remanent population represented the cells with a round nuclei. All the results are presented as percentage of the population with a given nuclei shape. For the Lamin A and Lamin B measurements, both, the Inner and Outer area of the nuclei were considered. Whole Nuclei were segmented based on the Hoechst channel. The outer region of the nucleus or nuclear envelope area, was defined by the maximum antibody intensity in the nuclear membrane (Figure S9K). For the Phospho-Lamin A to Lamin A ratio determination, the intensity of the P-Lamin in the Inner region or nucleoplasm was divided by the intensity of Lamin A in the outer region of the nuclei or nuclear membrane. YAP, TAZ or YAP/TAZ ratios were calculated as the log10 of the mean Inner nuclear intensity/mean perinuclear region intensity per cell. YAP and TAZ ratios were corrected for cell number on mock-transfected cells by using a linear regression analysis (Sero et al., 2015). In each experiment, different numbers of mock transfected cells were plated to cover the range of densities of siRNA transfected cells. The observed value for the log10 TAZ nuc/cytop ratio or log10 YAP nuc/cytop ratio was measured and corrected by using the linear regression equation and the observed minus predicted value was plotted in all the figures of this manuscript.

<u>Organoid segmentation</u>, the **‘**find image’ region module was used to define the area of the core of the organoids based on the actin channel and split into objects as a population named ‘colonies’. Then, within that population, cells were found based on the Hoechst channel. This feature was used to classify the colonies population according to the number of cells per colony, in Single cells (One nuclei per colony) or Acini, as two or more nuclei per colony. Further characterization of organoids was carried out exclusively in the Acini population: fibronectin mean intensity of the Acini body area was measured and normalized to the mean area of the Acini. Actin protrusions or spikes were determined by using the Find neurites modulus and based on the actin channel. To perform all the analysis in 3D, the maximum projection was used.

<u>For the bottom to top invasion assays</u>, nuclei were segmented as above and cell cores were segmented on the actin channel. A contrast threshold for the Inner nuclei was chosen for every experiment (commonly 0,50) to select only cells that were really in the selected plane to avoid quantifying the same cell in several planes. A Second step to filter cells was performed by excluding the cells were the mean of the actin Intensity in the perinuclear area was lower than 150 (Figure S10B II). The invasion index was calculated as the sum of the number of cells at 30 μm plus 60 μm and divided by the total number of cells (0 μm + 30 μm + 60 μm).

### Sample preparation for MS analysis

Cells were transfected with the siRNAs as stated above and/or treated with DMSO or RBL at 40uM at day cero and collected 48h later by trypsinization. After resuspension in growth media and centrifugation, media was removed and 1mL of cold PBS was added. Then one million of viable cells per condition was transferred to low binding mass spectrometry tubes and washed 2 x with cold PBS to a final pellet that was flash frozen with 70% ethanol and dry ice. After that, cell pellets were dissolved in 1% sodium deoxycholate (SDC), 100mM triethylammonium bicarbonate (TEAB), 10% isopropanol, 50mM NaCl and Halt protease and phosphatase inhibitor cocktail (100X) (Thermo, #78442) with pulsed probe sonication for 15 sec and boiling at 90 °C for 5 min followed by re-sonication for 5 sec. Protein concentration was measured with the Coomassie Plus Bradford Protein Assay (Pierce). Aliquots of 100 μg of total protein were reduced with 5 mM tris-2-carboxyethyl phosphine (TCEP) for 1 h at 60 °C and alkylated with 10 mM Iodoacetamide (IAA) for 30 min in dark followed by trypsin digestion overnight (Pierce, 75 ng/μL). The peptides were labelled with the TMT-10plex multiplex reagents (Thermo) according to manufacturer instructions and were combined in equal amounts into a single tube. The peptide mixture was fractionated with high pH Reversed-Phase (RP) chromatography using the XBridge C18 column (2.1 x 150 mm, 3.5 μm, Waters) on a Dionex Ultimate 3000 HPLC system over a 1% gradient in 35 min. Mobile phase A was 0.1% ammonium hydroxide and mobile phase B was 0.1% ammonium hydroxide in acetonitrile. Phosphopeptide enrichment was performed on the fractions with the High-Select Fe-NTA kit (Thermo) according to manufacturer instructions. The LC-MS analysis was performed on the Dionex Ultimate 3000 system coupled with the Orbitrap Fusion Lumos Mass Spectrometer (Thermo) using the EASY-Spray C18 capillary column (75 μm × 50 cm, 2 μm) over a 90 min gradient. MS spectra were collected with mass resolution of 120k and precursors were targeted for CID fragmentation in the top speed mode. CID collision energy was set at 35% with IT 50 ms. Peptides were quantified at the MS3 level with HCD fragmentation (collision energy 65%, 50k resolution) of the top 5 most abundant CID fragments isolated with Synchronous Precursor Selection (SPS) and with 105 ms IT. Targeted precursors were dynamically excluded for further activation for 45 seconds. For the phosphopeptide analysis, MS2 spectra were collected with HCD fragmentation (CE 38%) and 50k resolution. The Sequest HT engine in Proteome Discoverer 2.2 (Thermo) was used to search the raw mass spectra against reviewed UniProt human proteins. The precursor mass tolerance was set at 20 ppm and the fragment ion mass tolerance was 0.5 Da for CID and 0.02 Da for HCD spectra. TMT6plex at N-terminus/K and Carbamidomethyl at C were defined as static modifications. Dynamic modifications were oxidation of M and Deamidation of N/Q as well as phosphorylation of S/T/Y for the phosphoproteome analysis. Peptide confidence was estimated with the Percolator node and peptide FDR was set at 0.01. The reporter ion quantifier node included a TMT10plex quantification method with an integration window tolerance of 15 ppm at the MS3 level. Only unique peptides were used for quantification, considering protein groups for peptide uniqueness. Peptides with average reporter signal-to-noise>3 were used for protein quantification.

### Subcellular fractionation and immunoblotting

The Subcellular Protein Fractionation Kit for cultured cells (ThermoFisher) was used to isolate the cytoplasm, membrane, soluble nuclear fraction, chromatin-bound and cytoskeletal protein extracts. MCF10A cells were harvested and washed by suspending the cell pellet with ice-cold PBS. Then, 1% NP-40/0.1% SDS lysis buffer (50 mM Tris 7.4, 150 mM NaCl, 1mM EDTA) with protease and phosphatase inhibitor cocktail (Thermo) was added. Protein concentrations of the subcellular protein fraction lysates were measured using Pierce BCA Protein Assay Kit (ThermoFisher). Samples were boiled for 10 minutes and the extracts were separated by SDS-polyacrylamide gel electrophoresis using precast 4–20% Precise protein gels (ThermoScientific). Separated proteins were transferred to PVDF-FL transfer membrane (Merck Millipore) at 40V for 90 min at 4 °C. PVDF membranes were blocked in 5% milk/TBS 10% glycerol + 0.05% tween20 (Sigma-Aldrich) (TBS/t) or 5% BSA/TBS 10% glycerol + 0.05% TBS/t for 1 hr at RT on a rolling shaker, then incubated in the appropriate primary antibodies: Mouse anti-GAPDH (Novus Biologics, 1:2000), Mouse anti-LMNA (Abcam, 1:500), Mouse anti-PCNA (Santa Cruz, 1:1000), Rabbit anti-KIF5B (Proteintech, 1:1000), Mouse anti-β Tubulin (Sigma, 1:1000) and Mouse anti-YAP/TAZ (Santa Cruz, 1:1000) overnight at 4 °C on a rolling shaker, washed 3 × 20 min in TBS/t before incubating for 1 hr at RT in secondary anti-rabbit HRP-linked antibody and/or anti-mouse HRP-linked antibody (Cell Signalling) on a rolling shaker. Membranes were washed 3 × 20 min in TBS/t, 1x in PBS for 5 min and then soaked in an enhanced luminol-based chemiluminescent substrate for detection of horseradish peroxidase (HRP) on immunoblots before visualising on the Azure c300 scanner. Western Blot analysis were performed using Image J and Excel.

### RNA extraction, PCR Array Human Epithelial to Mesenchymal Transition (EMT) and qRT–PCR

Total RNA was extracted using the standard phenol-chloroform procedures (Trizol) and the RNeasy kit (Qiagen). For the RT² Profiler™ PCR Array Human Epithelial to Mesenchymal Transition kit (EMT) (Qiagen), RNA integrity of all the samples was tested by using the Agilent RNA 6000 Nano kit (Agilent Technologies, Cheshire, UK) and measured with an Agilent Bioanalyser. The RNA integrity number (RIN) was calculated by using the Agilent 2100 software. Briefly, 400 ng of each RNA was used following supplier’s recommendations and carried out on a Quant 6 RT PCR (Applied Biosystems). Gene expression data was analyzed by excel and normalized to the mock transfected values. Gene expression numbers lower than 0.5 were considered as downregulated and genes with values greater than 1.3 were considered as upregulated. Heat maps were generated by using the Morpheus software (https://software.broadinstitute.org/morpheus). Panel of genes included in the EMT kit and results are showed in Table S3, RT² Profiler™ PCR Array Human EMT_Cat# PAHS-090Z_Data related to Fig3G_Gene expression of JAM3 siRNAs. The conventional Quantitative real time polymerase chain reaction (qRT-PCR) was performed using cDNA made from extracted RNA. Briefly, 250 ng of purified RNA was converted to cDNA by using the High-Capacity RNA-to-cDNA Kit (Applied Biosystems) according to the manufacturer’s protocol. qRT-PCR analysis was carried out using the SYBRGreen PCR Master Mix (Invitrogen) following supplier’s recommendations, on a 7300 AB system (Applied Biosystems) or a Quant 6 RT PCR (AppliedBiosystems). ΔΔCt quantification was performed on at least triplicate absolute measurements and normalized to GAPDH mRNA.

### Live RELA imaging

RELA was tagged at the endogenous C-terminus using CRISPR/CAS9 in RPE1 cells previously tagged with AAV mRuby-PCNA (Barr et al., 2017). RPE1 mRuby-PCNA RELA-GFP were seeded at 250 cells/well in 30µl DMEM in a 384 well plate (PerkinElmer). The following day, cells were transfected with non-targeting or JAM3 siRNA with Lipofectamine RNAiMAX (Thermofisher) according to the manufacturer’s protocol. 48 hr after siRNA transfection, cells were imaged at 10 min intervals for 2 hours then imaged for a further 5 hr following TNF (R&D Systems) treatment (0, 0.1 or 10 ng/ml). Imaging was carried out with a 20x objective on the PerkinElmer Opera confocal microscope with an environmental control chamber set to 80 % humidity, 5 % CO2 and 37 °C. Data were analysed using Columbus software (PerkinElmer).

### Statistics and Reproducibility

All data generated in this manuscript was derived from at least two independent experiments and are presented as means ± SEM. Comparisons among groups were performed using GraphPad Prism version 6.00 and 7.00, GraphPad Software, La Jolla California USA, www.graphpad.com. Analysis of regression lines and *R^2^* values (Pearson’s correlation) and qRT-PCR analysis were performed with Excel. Statistical details of experiments can be found in the legends.

Survival analysis was performed with breast cancer–specific 10-year survival data. Kaplan-Meier estimator was used for patient stratification where log-rank test was used for testing difference among groups. Cox proportional hazards regression model was fitted, and 95% confidence intervals were computed to determine the prognostic values, where log-rank test with *P* < 0.05 was considered significant. Correlation was computed with Pearson correlation and q-values computed using False Discovery Rate (FDR) correction using R package fdrtool in the situation of multiple correlation analysis. Test for significant differences among groups of a single variable was carried out using ANOVA. Tests for trend of a continuous variable among groups were performed with Jonckheere’s Trend test for a null hypothesis that a variable does not follow a monotone trend set by the groups.

### The PRESTIGE framework for network analysis

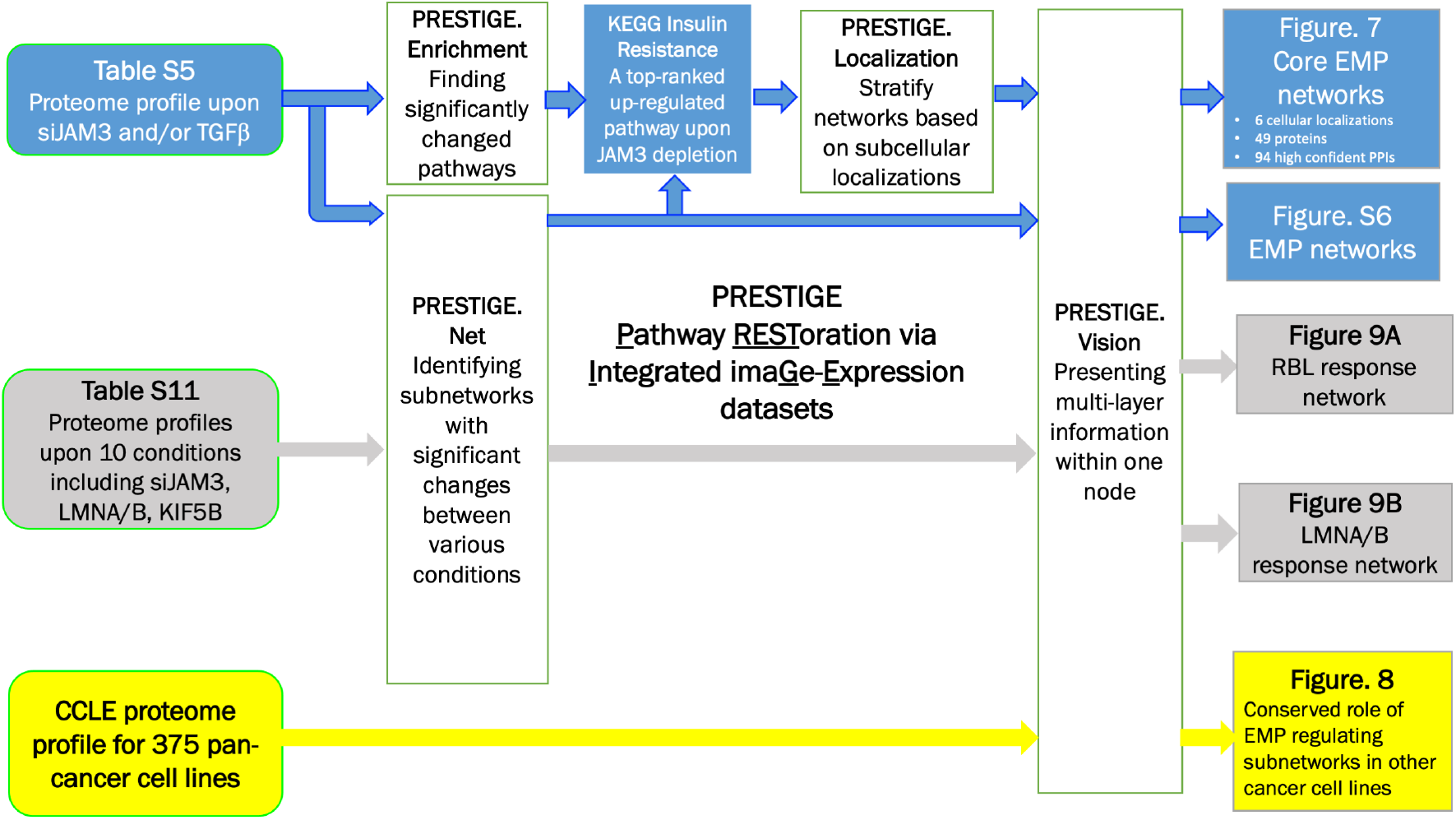

The customized systems biology framework of PRESTIGE (**<u>P</u>**athway **<u>REST</u>**oration via Integrated ima**<u>G</u>**e-**<u>E</u>**xpression datasets) was designed and applied to three proteomics datasets from MCF10A breast cancer cell lines, i.e.

Effect of JAM3 depletion and TGFβ treatment, as presented in Table S5, with total protein and phosphorylation data from four conditions:

1. untreated wildtype cells denoted as Wildtype,
2. depletion of JAM3 (denoted as siJAM3),
3. TGFβ treatment, and
4. combined treatment (denoted as siJAM3+TGFβ).

Proteomic profile as presented in Table S12, with total protein and phosphorylation data from ten conditions:

1. untreated wildtype cells denoted as control
2. JAM3 (SMARTpool)
3. JAM3 (OTP5 version)
4. 4) LMNA
5. 5) LMNB
6. 6) KIF5B
7. 7) SYNE4
8. KIF5B, SYNE4 and JAM3 (SMARTpool)
9. control+KIF5B inhibitor (RBL)
10. JAM3 (SMARTpool)+KIF5B inhibitor (RBL)

### Recording protein responses to various perturbations of interest

Total protein intensities for N=8,268 proteins were obtained from the aforementioned four conditions. After normalization and scaling, each protein is be described by a 4-tuple intensity vector 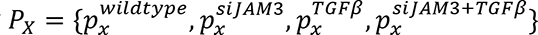, and a 5-tuple fold change vector

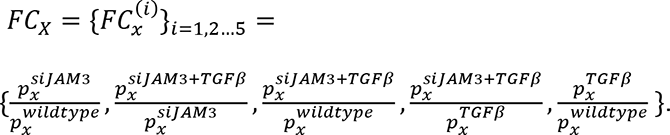

Here 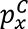 indicates the average value of total protein levels for x measured from samples in condition *C*. Also, FC^(1)^ and FC^(5)^ illustrate the effect of single perturbation siJAM3 and TGFβ, respectively, when compared to untreated wildtype; FC^(2)^ and FC^(4)^ indicate the effect of siJAM3+TGFβ combination as compared to siJAM3 and TGFβ, respectively; and FC^(3)^ represents the combination effect as compared to untreated wildtype.

### Linking JAM3 to PPI networks among proteins significantly increased by siJAM3 and/or TGFβ

We collected 234 proteins significantly increased by siJAM3 and/or TGFβ perturbations, including:
99 proteins with FC^(1)^ ≥1.2 and FC^(2)^ ≥1.1;
68 proteins with FC^(5)^ ≥1.2 and FC^(4)^ ≥1.1;
44 proteins with FC^(i)^ ≥2.5 in any *i* ∈ [1,5];
22 proteins with 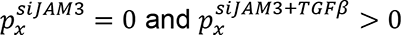;
10 proteins with FC^(i)^ ≥1.5, FC^(j)^ ≤ 5/6 and FC^(k)^ ≤ 10/11, (*j*, *k*) ∈ {(1,2), (5,4)};

with 9 proteins belonging to more than one of these 5 groups.

These proteins were combined with YAP1, WWTR1, KIF5B, and SYNE4 and served as input for PPI query in STRING database (Ver. 11.0), and there are 228 PPIs with combined confidence score≥0.4 amongst 150 of these 238 proteins. INS (Uniprot Accession P01308, Insulin) was involved in 22 of these 228 PPIs and was by far the protein with the most PPIs within this group, as no other proteins had more than 15 PPIs.

### Identifying member of KEGG Insulin Resistance pathway that are increased by siJAM3

Given the significantly high number of PPIs involving INS in the above analysis, Wilcoxon enrichment analysis was run on 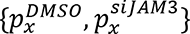 for all 8,267 available proteins using ConsensusPathDB (Release 34) to identify pathways significantly up-regulated following siJAM3. Among all the pathways containing INS, KEGG_Insulin_Resistance (HSA04931) had an enrichment p-value of 0.00948, q-value of 0.0853, only 8 of 65 member proteins for HSA04931 had FC^(1)^ ≤0.85, comparing to 26 with FC^(1)^ ≥1.2, including INS, IRS1, IRS2, as well as RELA and IKBKB.

### Generating and visualizing the final networks model for JAM3’s role in regulating EMT

The result from Wilcoxon enrichment analysis indicates that siJAM3 strongly up-regulated insulin and NFκB related pathways and prompt us to combine these 26 up-regulated proteins with the 238 proteins identified from the previous section and re-run the PPI query. Overall, 197 PPIs with combined confidence score≥0.7 were identified from 108 of 260 proteins identified above. INS had 17 PPIs and remained the most connected protein in the network, while PTEN, FOXO1, IRS1, IKBKB, and AKT1 all had more than 10 PPIs with confidence score≥0.7.

The EnhancedGraphics plugin (Ver. 1.5.4) was used in Cytoscape (Ver. 3.8.2) to visualize the resulted networks. Each protein was represented by a circle node, and each circle was evenly split into five sectors with central angle of 72^°^. Starting from the 6 o’clock direction and going clockwise within each circle, the color of the five sectors were proportional to the log2 transformation of FC^(1)^, FC^(2)^ …FC^(5)^, respectively, thus leaving the 12 o’clock direction covered by the sector representing the effect for siJAM3+ TGFβ, i.e.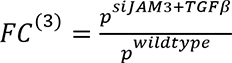. For each sector, deeper red color corresponds to stronger up-regulation while deeper blue color corresponds to stronger down-regulation. For each circle, the size is proportional to the number of PPIs involving the corresponding protein, and member proteins for KEGG_Insulin_Resistance (HSA04931) were highlighted using a red boundary round the circle. Qualified (combined confidence score≥0.7) PPIs between two proteins were represented by one single edge connecting two circles, and the thickness of an edge were proportional to the combined confidence score for the corresponding PPI. The final figure was shown as Figure S6.

### Visualization of JAM3-Insulin Resistance-NFκB interaction networks as stratified by subcellular localization

Uniprot Accession numbers (release 2021-01) for the 260 proteins shown in the EMP network were used to query InnateDB (Ver. 5.4), and 218 proteins were assigned to one of the five subcellular locations, i.e. extracellular matrix, cell surface, plasma membrane, cytoplasm, and nucleus. Within the EMP network, all interaction partners with the 25 members of KEGG Insulin Resistance pathway (HSA04931) were collected and a core EMP network with 94 PPIs amongst 48 proteins stratified in five subcellular locations was formed as Figure 7. The nodes had same colour pattern as in Figure S6, with 25 members of HSA0493 highlighted using pink squares behind the round nodes. The five sub-cellular localizations were illustrated using different colours.

### Benchmark the behaviours of siJAM3/TGFβ induced pathways and networks in pan-cancer cell lines using proteomics data from Cancer Cell Line Encyclopaedia (CCLE)

Although the EMT regulating networks presented in Figures S6 and 7 as well as were identified using proteomics data from MCF10A cell lines, such networks had the potential to bear conserved functions in a wide range of pan-cancer cell lines. To define the extent where the EMP regulation function of our networks were conserved, the dataset for normalized total protein abundances for 12,490 unique proteins across 375 unique CCLE cell lines was downloaded from this link: https://gygi.hms.harvard.edu/data/ccle/protein_quant_current_normalized.csv.gz

In this dataset, the total protein abundance for each protein in each cell line was represented by a z-score. For each cell line L, the average z-scores for all available members from EMP network, core EMT network and the 25 members in Table S6 belonging to KEGG insulin resistance pathway (HSA04931) were calculated and summarized in a vector M^L^, defined by [mean_EMP^L^, mean_Core_EMP^L^, mean_IR^L^]. The following two conditions were applied to search for cell lines where features in M^L^ are consistent with significantly changed canonical EMT marker VIM, specifically:

26 cell lines had z-score for VIM≥1.5, and all three features in M^L^≥0,
30 cell lines had z-score for VIM≤-1.5, and all three features in M^L^≥0.

Hierarchical clustering was carried out using the CCLE profile for all available Table S6 members across these 56 cell lines and the resulted heat map was shown as Figure 8A. PCCs were used as similarity measurements between cell lines (columns) and proteins (rows), and Morpheus from Broad Institute was used for clustering and creation of heatmaps.

In a 3D space defined by [mean_EMP, mean_Core_EMP, mean_IR], the 56 cell lines with positive vs. negative VIM values were separated in to two non-overlap groups, as shown in Figure 8B. The following silhouette widths were calculated on vector M^L^ for all 56 cell lines and used for cluster validation:

For cell line *L* ∈ *F*_*I*_, where clusters 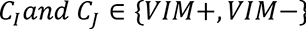, where *d*(*L*,*N*) is the Euclidean distance between three tuple vectors M^L^ and M^N^, and

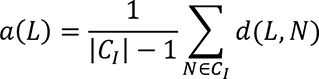

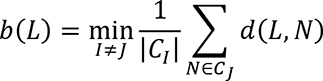

Silhouette width for cell line L is thus defined as:

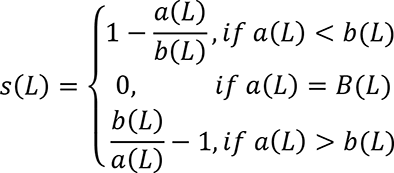

All but one cell lines had positive silhouette width and 52 out of 56 cell lines had silhouette width larger than 0.5. MATLAB R2020a Update 5 (9.8.0.1451342) maci64 was used on a MacPro workstation for the calculation of silhouette width.

Regarding the specific case where lowered JAM3 levels corresponds with accelerated EMT processes, 171 cell lines in the CCLE proteomics dataset had VIM>CDH1 while also having detectable JAM3, and 99 of these 171 cell lines had negative z-scores for CDH1, which shows that the depletion of JAM3 is non-exclusive with the depletion of CDH1.

### Define the subnetwork responding to KIF5B

Student’s t-tests were applied to identify significantly changed total protein or phosphorylation sites between the two groups of conditions:

Group A: 2) JAM3 (SMARTpool) and 3) JAM3 (OTP5 version)
Group B: 6) KIF5B, 9) control+KIF5B inhibitor (RBL) and 10) JAM3

(SMARTpool)+KIF5B inhibitor (RBL).

As summarized in Table S12, the scaled abundance of 190 phosphorylation sites and 105 total proteins had p-value≤0.05 from student’s t-test and more than 2-fold changes in either direction, including 96 phosphorylation sites and 44 total proteins that are up-regulated upon KIF5B treatments. These significantly changed sites and proteins are associated with 260 unique proteins and according to STRING database, 119 of these 260 proteins involved in 153 high confidence PPIs (confidence score≥0.7). Each protein is represented by one round node and each significantly changed phosphorylation site is represented by one ring around the node for its master protein. The colour of a node of ring is proportional to the −log10 transformation of p-values from the pooled comparisons on KIF5B vs. JAM3 depletions, with red colour denote up-regulation upon KIF5B treatment. The size of a node is proportional to the number of PPIs it involved in the figure. 12 of the 115 proteins in this figure are the members of the EMP network and are highlighted by a pink square in the back of the round node. Same as in Figure S6 and Figure 7, the EnhancedGraphics plugin (Ver. 1.5.4) was used in Cytoscape (Ver. 3.8.2) to visualize the resulted networks.

### Define the subnetwork responding to LMNA/LMNB and interacting with the core EMT network

Student’s t-test was applied to identify significantly changed total protein or phosphorylation sites between the two groups of conditions:

Group A: 1) control, 6) KIF5B, and 7) SYNE4
Group B: 4) LMNA, 5) LMNB, 8) KIF5B, SYNE4 and JAM3 (SMARTpool), 9)

control+KIF5B inhibitor (RBL), and 10) JAM3 (SMARTpool)+KIF5B inhibitor (RBL).

337 total proteins and 405 phosphorylation sites were significantly up-regulated in LMNA/LMNB cells, which corresponds to 623 unique proteins. According to STRING database, 94 of 623 LMNA/LMNB proteins interacted with 40/48 members of the core EMT network. There are 191 high confidence (confidence score≥0.7) PPIs involving one LMNA/LMNB protein and one member of the core EMT network.

Cytoscape (Ver. 3.8.2) was used to visualize the resulted networks. The 94 proteins highlighted above was represented by purple circles, and 3 of the 94 proteins had a red square in the back of their purple circles because they are also members of core EMT network. Other members of the core EMT networks are denoted by red circles. Overall, these 94 proteins involved in 191 high confidence PPIs with 40/48 members of core EMT networks (red circles if not within the group of 94). High confidence PPIs within two proteins of the same colour were also included to make a network of 295 PPIs amongst 136 unique proteins (91 purple circles, 3 purple circles with red squares, 37 red circles interacting with purple circles, 5 red circles only interacting within this group of 5 and not with purple circles). The size of a node is proportional to the number of PPIs it involved in the figure.

---

**Figure S1.**
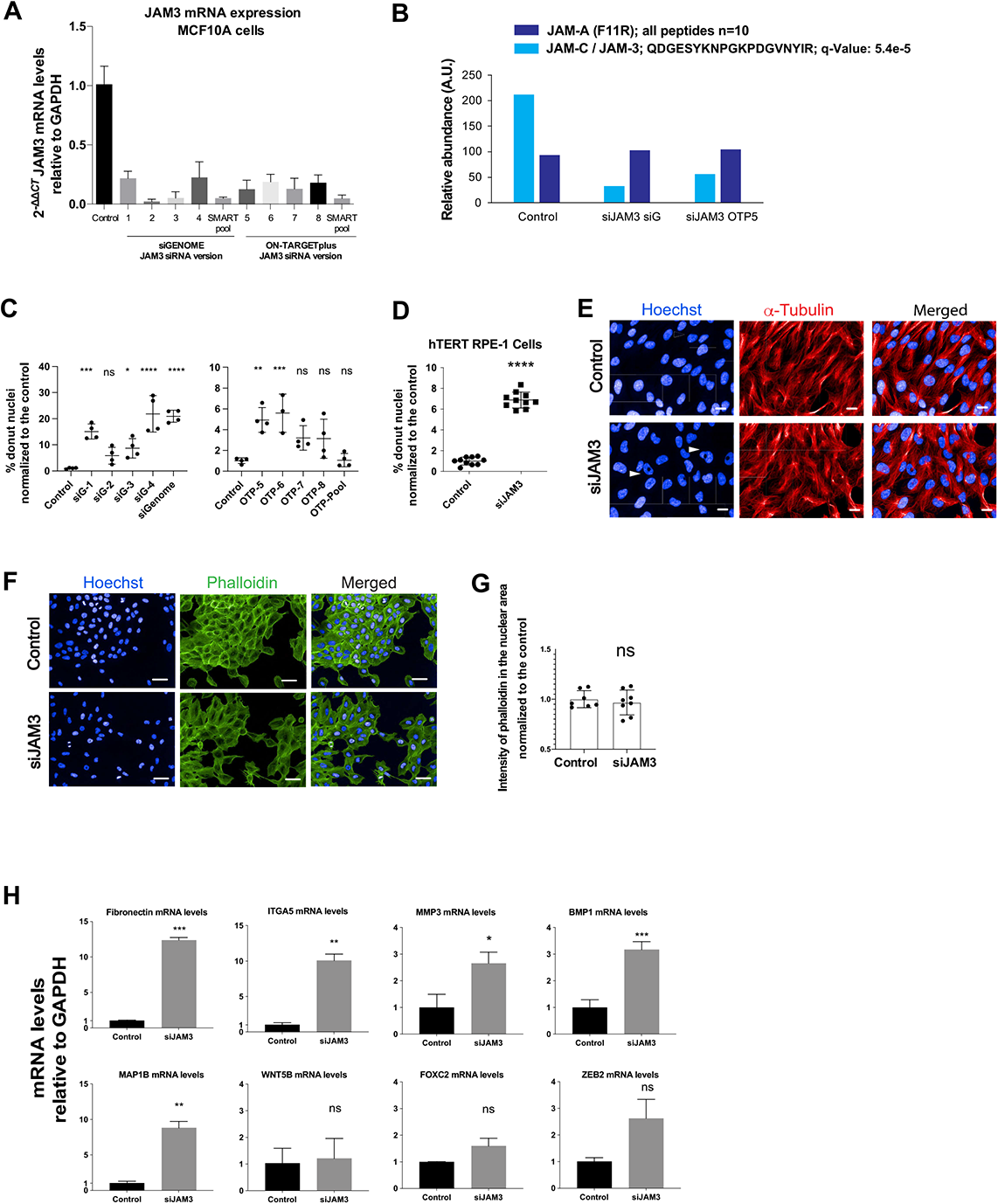
On-target effects of JAM3 siRNA. (A) JAM3 mRNA levels as determined by qRT-PCR following depletion by single or pooled siRNAs. Data was normalized to GAPDH mRNA and control (wild type). Mean +/- SD (n= 4 wells/condition). (B) Detection of JAM3/JAM-C and JAM-A protein abundance by Proteomics in Mock transfected MCF10A cells, siGenome (siG) or On Target plus pool 5 targeting JAM3. (C) Percentage of cells with nuclear donuts in Control and JAM3 depleted cells by using single and pooled siRNAs targeting JAM3. Both graphs show a single experiment (n > 4.000 cells). Comparisons among groups were performed by one-way ANOVA (Newman-Keuls multiple comparison test) with Prism6 software (∗p < 0.05, ∗∗p < 0.01, ∗∗∗p < 0.001, ∗∗∗∗p < 0.0001). (D) Percentage of cells with donut shape in Control or JAM3 depleted hTERT1 RPE-1 cells. Data shows the results of 2 different experiments (n > 7.000 cells). t student was performed with Prism6 software (∗∗∗p < 0.001). (E) Representative images of the nuclei (blue, Hoechst) and microtubules (red, α-tubulin) in Control and JAM-3 depleted hTERT1 RPE-1 cells. Scale bars, 50um. (F) Representative images of the nuclei (blue, Hoechst) and phalloidin (green, actin) of MCF10A cells mock transfected or transfected with JAM3 RNAi. Scale bars, 50um. (G) Quantification of the total actin intensity in the nuclear area and the texture of the actin fibers in the whole cell area in MCF10A cells mock transfected and JAM3 depleted. Graphs show the data of one experiment (n > 5.000 cells). t student was performed with Prism6 software. (H) Confirmation of gene expression with individual primers. Relative mRNA levels of Microtubule Associated Protein 1B (MAP1B), Transcription factors such as FOXC2 or the ZEB2 and genes related to ECM organization such as Fibronectin 1 (FN1), Integrin Subunit Alpha 5 (ITGA5), Stromelysin 1 (MMP3) or Procollagen C-Endopeptidase (BMP1) were normalized to GAPDH for wild-type and JAM3 depleted cells. Mean ± SD (n = 4 replicates/condition). (∗p < 0.05, ∗∗p < 0.01).

**Figure S2.**
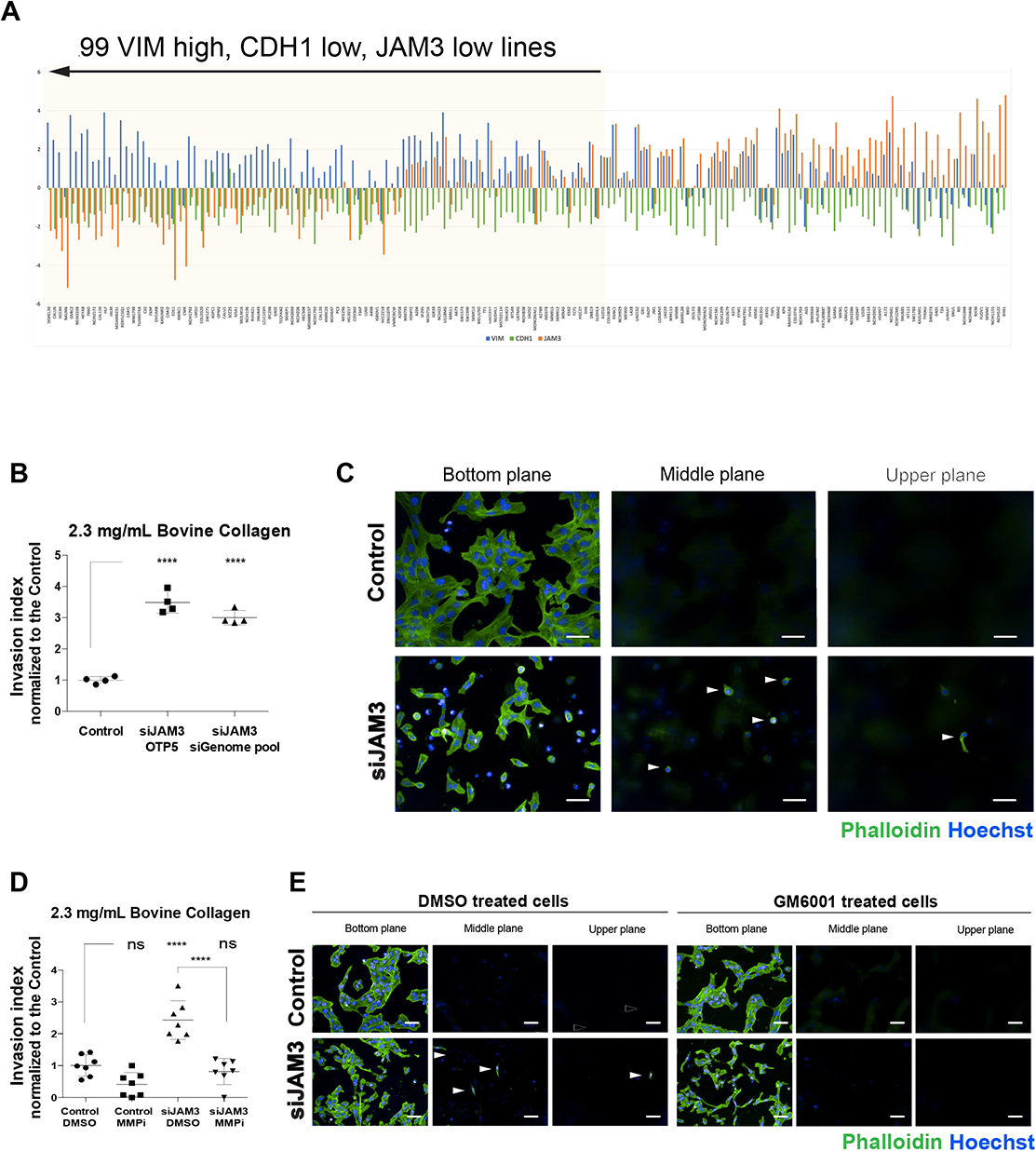
Low JAM3 levels correlate with mesenchymal cancer and result in invasiveness. (A) Z-scores for VIM (blue), CDH1 (green) and JAM3 (orange) for 171 pan-cancer cell lines based on proteomics data from Cancer Cell Line Encyclopaedia (CCLE). All cell lines had higher z-score for VIM than CDH1. The cell lines were sorted from left to right in descending order of the differences between the VIM and JAM3 Z-scores. The first 99 cell lines on the left have higher z-score for VIM than JAM3. (B,E) MCF10A cells were allowed to invade “upward” into a Bovine Collagen I matrix, fixed, and labelled with Phalloidin (green) and Hoescht (blue), and then imaged at three different planes. (B) Invasion index of control and JAM3 depleted MCF10A cells. Representative data from 2 different experiments (n > 5.000 cells). Comparisons among groups were performed by one-way ANOVA (Newman-Keuls multiple comparison test; ∗p < 0.05, ∗∗p < 0.01, ∗∗∗p < 0.001, ∗∗∗∗p < 0.0001). (C) Representative images of cells in the 3 different planes imaged at 0, 30 and 60um. Scale bars, 50um (D) Invasion index of control and JAM3 depleted MCF10A cells treated with DMSO or MMP inhibitor GM6001. Graph shows the data of two different experiments (n > 5.000 cells). Comparisons among groups were performed by one-way ANOVA (Newman-Keuls multiple comparison test; ∗p < 0.05, ∗∗p < 0.01, ∗∗∗p < 0.001, ∗∗∗∗p < 0.0001). (E) Representative images of cells in the 3 different invasion planes imaged at 0, 30 and 60um. Scale bars, 50um.

**Figure S3.**
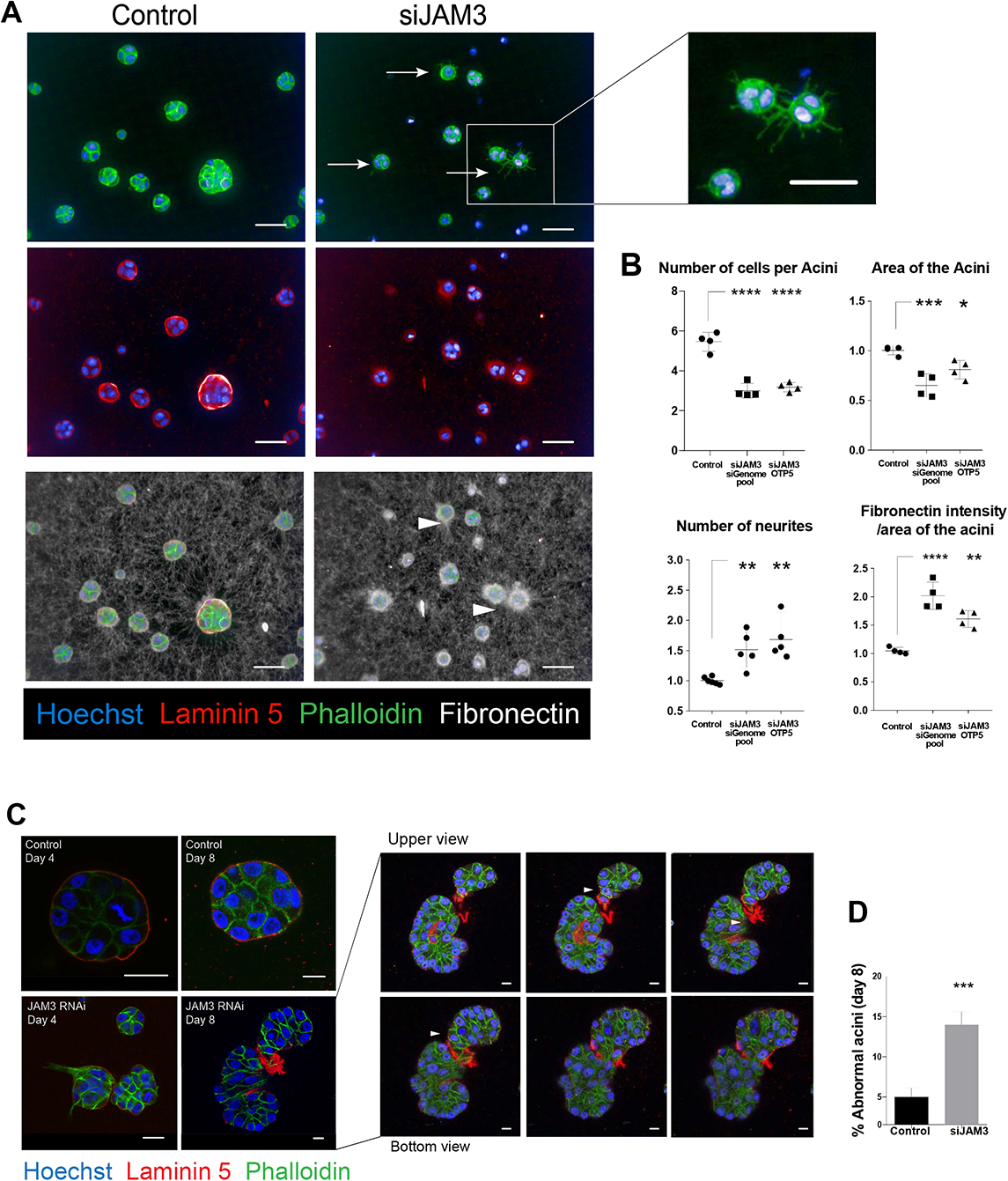
JAM3 is required for the morphogenesis of epithelial morphogenesis in 3D. (A,B) MCF10A Control or JAM3-depleted cells by 2 different siRNAs (OTP or siGENOME) growing in a Matrigel:Collagen thick matrix, allowed to form acini for 2 days, fixed and inmunostained. (A) Maximum projection of representative images of acini labelled with Hoechst (blue) and inmunostained for phalloidin (green), Laminin5 (red) and Fibronectin (white). Scale bars, 50um. (B) Graphs showing the morphology features of the 3 conditions. The majority of the structures formed had 2 or more nuclei, hence considered as Acini. Area was normalized to the Control. F-actin protrusions of the acini were described as Number of neurites and normalized to the control. Fibronectin intensity of the acini was expressed as the mean fibronectin intensity of the whole acini divided by the area of the acini and normalized to the Control. Data shows the result of one experiment where 4 wells per condition were analysed (n > 200 acini per condition). Comparisons among groups were performed by one-way ANOVA (Dunnett’s multiple comparison test) with Prism6 software (∗p < 0.05, ∗∗p < 0.01, ∗∗∗p < 0.001, ∗∗∗∗p < 0.0001). (C) Confocal sections of Control or JAM3-depleted MCF10A acini in a Matrigel:Collagen mix, 4 days or 8 days after transfection and labelled with Phalloidin (green), Laminin5 (red) and Hoeschst (blue). Scale bars indicated in the figures. (D) Visual quantification of the number of structures forming normal (round) or abnormal (irregular shape, multiple lobules or protrusive acini) acini at day 8 and expressed as a percentage (n = 100 acini). t student was performed with Prism6 software (∗p < 0.05, ∗∗p < 0.01, ∗∗∗p < 0.001, ∗∗∗∗p < 0.0001).

**Figure S4.**
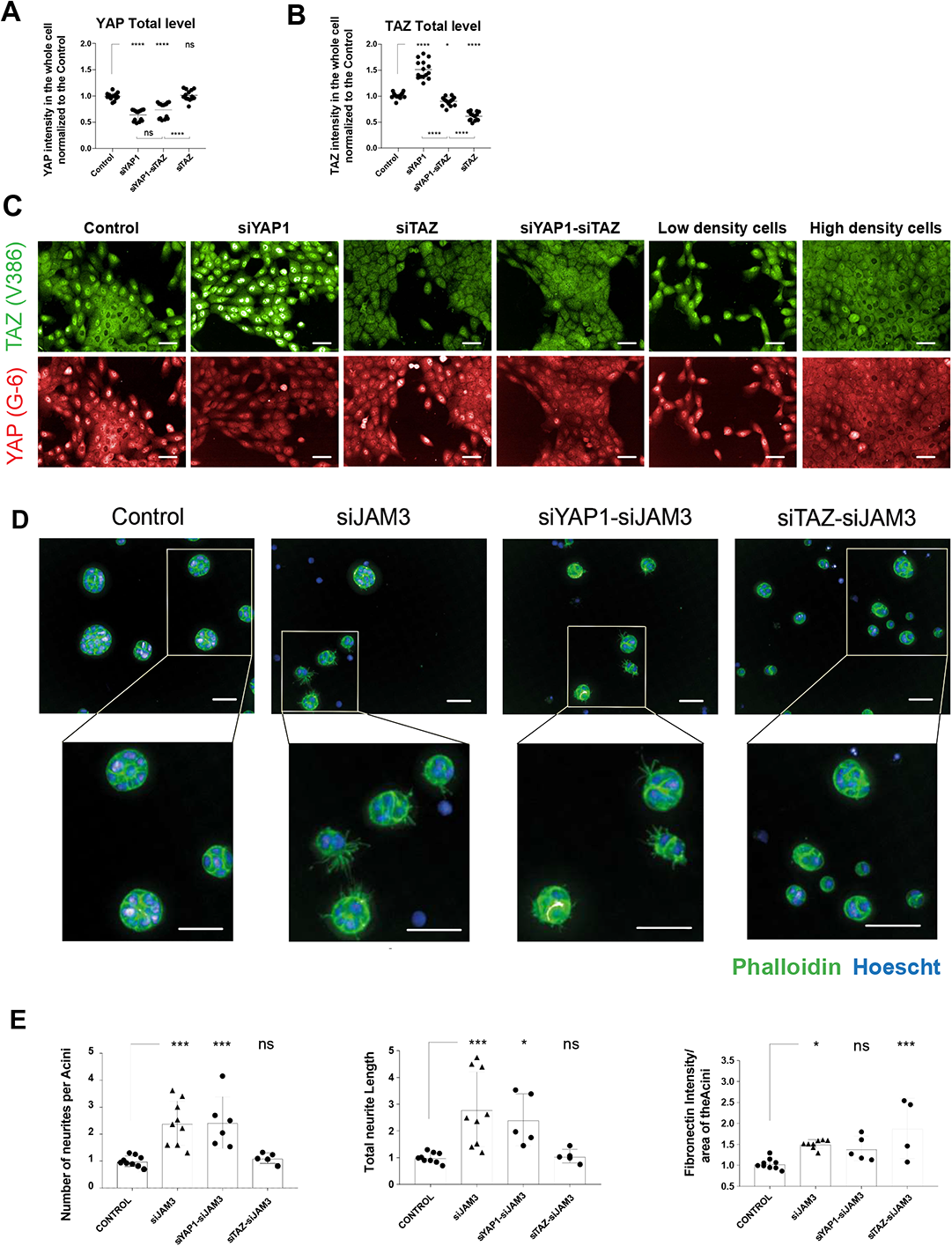
TAZ is required for EMT in 3D. (A) Quantification of total YAP intensity (whole cell) and (B) total TAZ intensity (whole cell) in Wild-type, YAP, TAZ and (YAP+TAZ) MCF10A depleted cells using antibodies specific for YAP and TAZ (n > 15.000 cells). Data shows the results of 3 different experiments. Comparisons among groups were performed by one-way ANOVA (Newman-Keuls multiple comparison test) with Prism6 software (∗p < 0.05, ∗∗p < 0.01, ∗∗∗p < 0.001, ∗∗∗∗p < 0.0001). (C) Representative images of Wild-type cells, treated with RNAi targeting YAP, TAZ or (YAP+TAZ) or seeded at high or low density and co-stained with TAZ (green) and YAP (red). Scale bars, 50um. (D,E) MCF10A Control or JAM3-depleted cells alone or in combination with YAP or TAZ siRNAs were plated in a Matrigel:Collagen thick matrix, allowed to form acini for 2 days, fixed and stained. (D) Maximum projection of representative images of acini labelled with Hoechst (blue) and phalloidin (green). Scale bars, 50um. (E) Graphs showing the morphology features of the 4 conditions. F-actin protrusions of the acini were described as Number of neurites per acini and also their length was measured and normalized to the control. Fibronectin intensity of the acini was expressed as the mean fibronectin intensity of the whole acini divided by the area of the acini and normalized to the Control. Data shows the result of two independent experiments where at least 2 wells per condition were analysed (n > 150 acini per condition). Comparisons among groups were performed by one-way ANOVA (Dunnett’s multiple comparison test) with Prism6 software (∗p < 0.05, ∗∗p < 0.01, ∗∗∗p < 0.001, ∗∗∗∗p < 0.0001).

**Figure S5.**
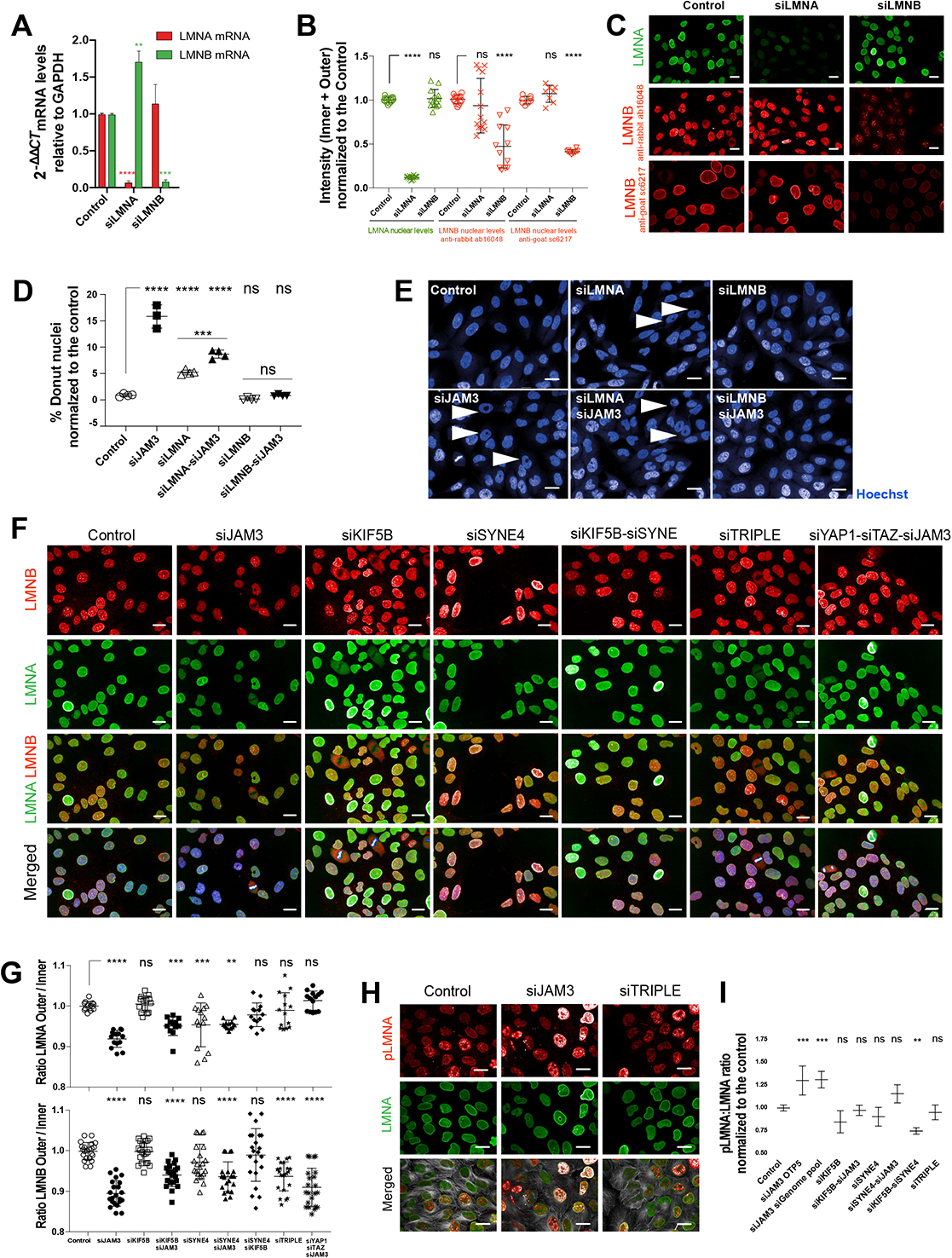
Nuclear morphology defects observed following JAM3 depletion are due to changes in the levels of LMNA and/or LMNB in the nuclear membrane. (A) Total mRNA levels of LMNA and LMNB normalized to GAPDH for wild-type cells. Mean ± SD (n = 3 replicates/condition) (∗p < 0.05, ∗∗p < 0.01). (B) Quantification of the total nuclear levels (Inner nuclear intensity + Outer nuclear Intensity) of either LMNA or LMNB in control MCF10A cells, or following depletion of LMNA or LMNB. Data shows the results of 3 different experiments (n > 15.000 cells). Comparisons among groups were performed by one-way ANOVA (Newman-Keuls multiple comparison test) with Prism6 software (∗p < 0.05, ∗∗p < 0.01, ∗∗∗p < 0.001, ∗∗∗∗p < 0.0001). (C) Representative images of Wild-type cells, treated with LMNA RNAi or LMNB RNAi and immunostained with LMNA antibody (green) or 2 different LMNB antibodies (red). Scale bars, 20um. (D,E) (D) Percentage of cells with nuclear donuts after LMNA or LMNB depletion alone or in combination with JAM3 (n > 15,000 cells). Data shows the results of 3 different experiments. Comparisons among groups were performed by one-way ANOVA (Šídák’s multiple comparisons test) with Prism6 software (∗p < 0.05, ∗∗p < 0.01, ∗∗∗p < 0.001, ∗∗∗∗p < 0.0001). (E) Nuclei of the cells labelled with Hoechst (blue). Arrows indicate donut nuclei. Scale bars, 20um. (F,G) MCF10A cells treated with different siRNAs and co-inmunostained with LMNA and LMNB. (F) Panel of images showing the staining of LMNA (green) and LMNB (red) after the RNAi treatments. Scale bars, 20 um. (G) Quantification of the nuclear LMNA and nuclear LMNB intensity normalized to the Control value. Data shows the results of 2 different experiments (n>15.000 cells). Comparisons among groups were performed by one-way ANOVA (Newman-Keuls multiple comparison test) with Prism6 software (∗p < 0.05, ∗∗p < 0.01, ∗∗∗p < 0.001, ∗∗∗∗p < 0.0001). (H,I) Cells were depleted of JAM3 alone, or in combination with KIF5B and/or SYNE4, and labelled with Hoechst, LMNA, phospho-LMNA and α-tubulin. (H) Images show cells stained with Hoechst (blue) and immunostained for LMNA (green), phospho-LMNA (red) and α-tubulin (grey). Scale bars, 20um. (I) Quantification of the ratio of phosphoLMNA in the inner nuclei or nucleoplasm/ LMNA in the Outer membrane after the RNAi treatments. Graphs shows the representative data of 2 different experiments (n > 5.000 cells). Comparisons among groups were performed by one-way ANOVA (Newman-Keuls multiple comparison test) with Prism6 software (∗p < 0.05, ∗∗p < 0.01, ∗∗∗p < 0.001, ∗∗∗∗p < 0.0001).

**Figure S6.**
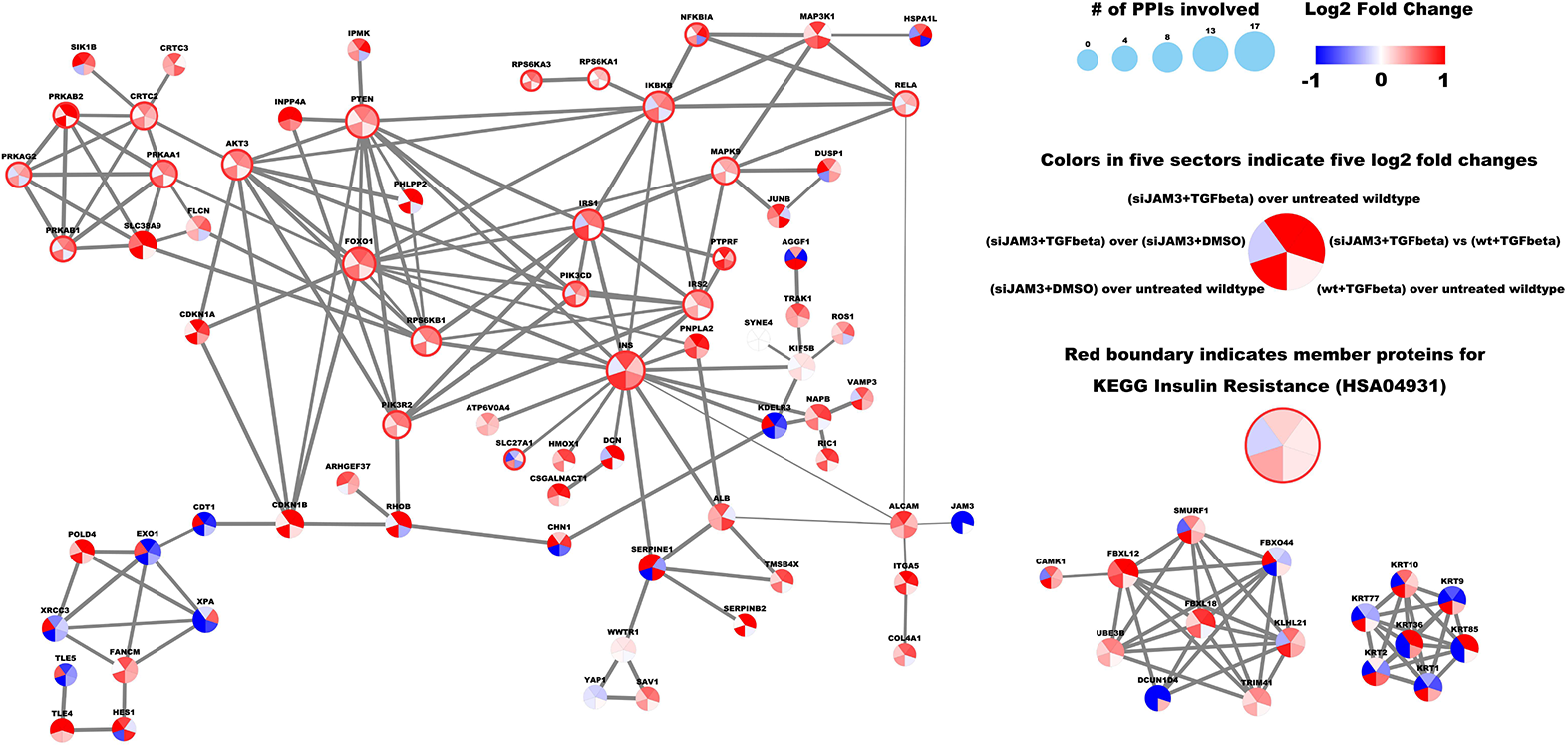
The three largest PPI connected neighborhood within the EMT network induced by JAM3 depletion and/or TGFβ treatment. There were 88 proteins involved in 186 PPIs and forming three PPI connected neighborhoods. Each protein was represented by one circle node that was evenly split into five sectors, and the color of the five sectors were proportional to the log2 transformation of fold changes from five different comparisons indicated in the legend section. All nodes showed significant up-regulation (red color as indicated in the legend) in at least one of the five comparisons, and more than half were up in at least three comparisons. Specifically, 22 of the 88 proteins belonged to KEGG Insulin Resistance pathway (HSA04931) and they were highlighted by red boundary around the node. One edge summarized all PPIs between the two connected nodes, as recorded in the STRING database, and the width of each edge was proportional to the overall confidence score given by STRING. The size of each node was proportional to the number of edges connected to it.

**Figure S7.**
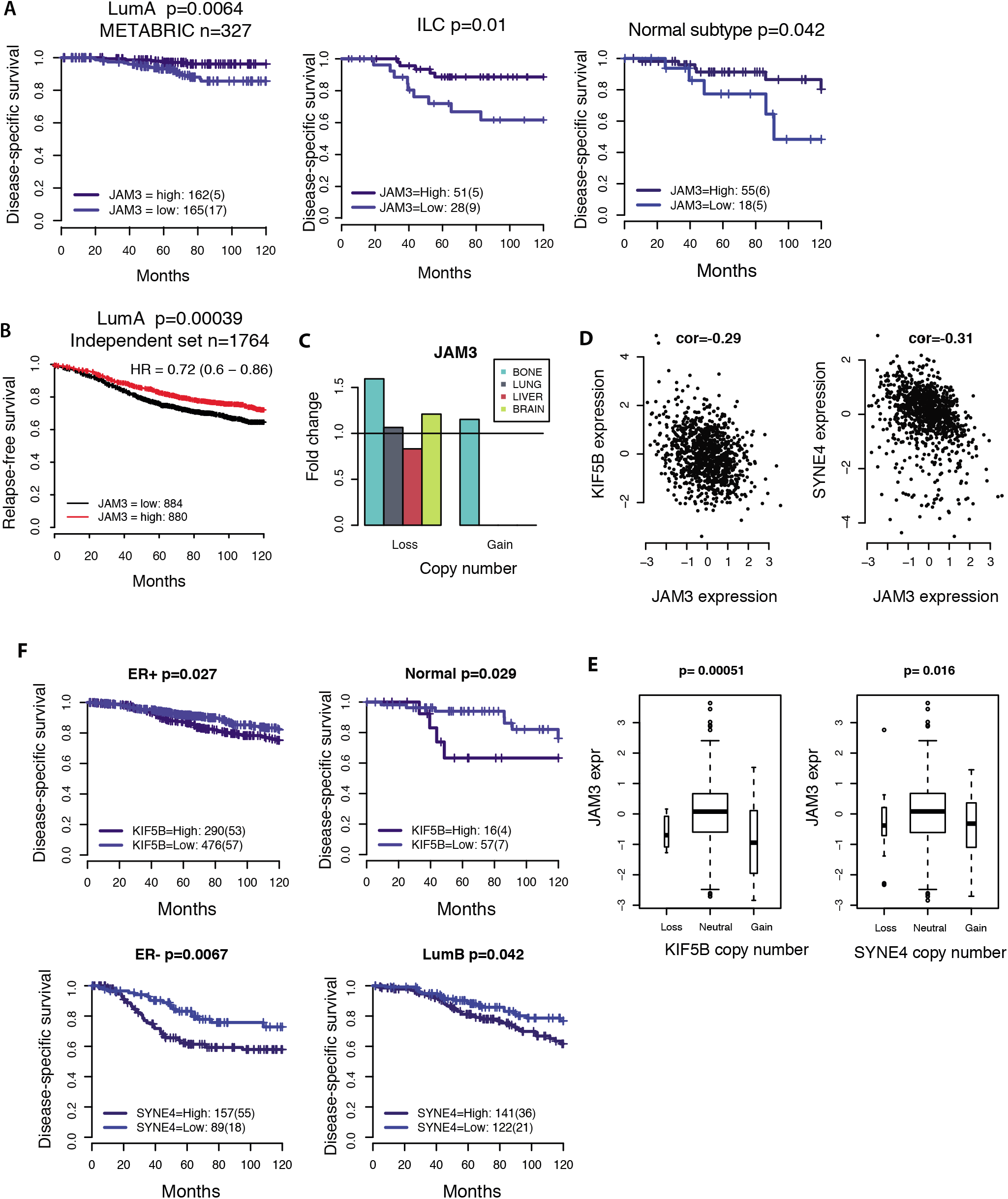
Regulation of FOXO3 nuclear translocation during EMP. (A-C) Wild-type or JAM3-depleted cells were non treated or TGFβ treated or TGFβ + RBL treated and co-stained with FOXO3a for image analysis. (A) Quantification of the nuclear:cytoplasmic FOXO3a ratio for all the conditions. Graph shows the representative data of 2 different experiments (n > 3.000 cells per condition). Comparisons among groups were performed by two-way ANOVA (Šídák’s multiple comparisons test) with Prism6 software (∗p < 0.05, ∗∗p < 0.01, ∗∗∗p < 0.001, ∗∗∗∗p < 0.0001). (B) Non treated control and JAM3 depleted cells were classified according to their nuclei shape and FOXO3a translocation was measured. Graph shows the representative data of 2 different experiments (n > 3.000 cells per condition). Comparisons among groups were performed by two-way ANOVA (Tukey’s multiple comparisons test) with Prism6 software (∗p < 0.05, ∗∗p < 0.01, ∗∗∗p < 0.001, ∗∗∗∗p < 0.0001). (C) Representative images of Hoechst (blue) and FOXO3a (green) for each condition where zoom images and arrows are showing the increase in the FOXO nuclear translocation upon TGFb and/or JAM3 RNAi treatments. Scale bars, 50um. (D) Model for regulation of FOXO3a by AKT during EMP.

**Figure S8. JAM-3 loss drives tumorigenesis and metastasis in breast cancer patients.**

(A-B) Kaplan-Meier curves illustrating the difference in disease-specific survival over 10 years according to JAM3 gene expression in different subtypes within METABRIC and in Luminal A subtypes of an independent cohort. ILC: Invasive Lobular Carcinoma. Within the luminal A subtype that had large number of samples, patients were spilt into equal-sized groups. For ILC and the Normal subtype, a best cut-off point was identified by testing different cut-offs. (C) Fold change in frequency of metastasis stratified by JAM3 copy number status. (D) Correlation between KIF5B/SYNE4 expression and JAM3 expression. (E) Boxplots illustrating the difference in JAM3 gene expression according to the copy number status of KIF5B and SYNE4. (F) Kaplan-Meier curves illustrating the difference in disease-specific survival over 10 years according to KIF5B/SYNE4 gene expression in different subtypes within METABRIC.

**Supplemental Table 1:** Nuclear shape feature names and definitions.

**Supplemental Table 2: Top 30 genes with the strongest correlation (positive or negative) between their gene expression and CNI.**

**Supplemental Table S4: Z-scores for VIM, CDH1 and JAM3 for 171 pan-cancer cell lines based on proteomics data from Cancer Cell Line Encyclopaedia (CCLE).**

**Supplemental Table S5: Scaled abundances for 8,267 total proteins and 12,529 phosphorylation sites across four conditions (untreated wildtype, TGFβ, JAM3 depletion and combined treatment of TGFβ+JAM3 depletion).**

**Supplemental Table S6: 260 proteins showed the trend of significant up-regulation upon depletion of JAM3 and/or TGFβ treatment.**

**Supplemental Table S7: Wilconxon enrichment analysis identified significantly up-regulated KEGG pathways upon depletion of JAM3 as compared to untreated wildtype MCF10A cells.**

**Supplemental Table S8: 3,247 of 12,529 phosphorylation sites showed significant changes across four conditions.**

**Supplemental Table S9: Overrepresentation analysis identified 69 pathways from KEGG or WikiPathways database that were enriched with master proteins for the 3,247 significantly changed phosphorylation sites across four conditions.**

**Supplemental Table S10: Benchmarking the behaviours of siJAM3/TGFβ induced pathways and networks in 56 pan-cancer cell lines using proteomics data from Cancer Cell Line Encyclopaedia (CCLE).**

**Supplemental Table S11: Scaled abundances for 9,227 total proteins and 18,975 phosphorylation sites across ten conditions.**

**Supplemental Table S12: Proteins and phosphorylation sites significantly changed after KIF5B related treatments.**

**Supplemental Table S13: 1,722 of 18,975 phosphorylation sites showed significant changes after KIF5B related treatments.**

**Supplemental Table S14: Overrepresentation analysis identified 68 pathways from KEGG or WikiPathways database that were enriched with master proteins for the 1,722 significantly changed phosphorylation sites after KIF5B related treatments.**

**Supplemental Table S15: Proteins and phosphorylation sites significantly changed after LMNA and LMNB related treatments.**

**Supplemental Table S16 : List of cell culture resources used in this study**

**Supplemental Table S17: Antibodies used in this study**

**Supplemental Table S18: siRNAs used in this study**

**Supplemental Table S19: qRT-PCR primers used in this study**

**Supplemental Table S20: Software used in this study**

